# Follicular fluid metabolome and cytokinome profiles in poor ovarian responders and the impact of dehydroepiandrosterone supplementation

**DOI:** 10.1101/2020.11.03.366047

**Authors:** Veronique Viardot-Foucault, Jieliang Zhou, Dexi Bi, Yoshihiko Takinami, Heng Hao Tan, Jerry.K.Y. Chan, Yie Hou Lee

**Author notes:** Emails: VVF; JLZ, DB, YT, HHT, JKYC, YHL. ***Address correspondence to:*** Jerry Chan MD, Ph.D, **E-mail:**, Lee Yie Hou Ph.D., **E-mail:**. These authors contributed equally (Veronique Viardot-Foucault, Jieliang Zhou).

## Abstract

Poor ovarian responders (POR) are women undergoing in-vitro fertilization who respond poorly to ovarian stimulation, resulting in the retrieval of lower number of oocytes, and subsequently lower pregnancy rates. The follicular fluid (FF) provides a crucial microenvironment for the proper development of follicles and oocytes. Conversely, dysregulated FF metabolome and cytokinome could have detrimental effects on oocytes in POR. Androgens such as dehydroepiandrosterone (DHEA) have been proposed to alter the POR follicular microenvironment but its effects on the FF metabolome and cytokine profiles is unknown. In this study, untargeted LC-MS/MS metabolomics was performed on FF of POR patients with DHEA supplementation (DHEA+) and without (DHEA-) in a randomized clinical trial (*N*=52). Untargeted metabolomics identified 118 FF metabolites of diverse chemistries, which included lipids, steroids, amino acids, hormones, among others. FF metabolomes were different between DHEA+ and DHEA- groups. Specifically, glycerophosphocholine, linoleic acid, progesterone, and valine were significantly lower in DHEA+ relative to DHEA-. Among cytokines, MCP1, IFNγ, LIF and VEGF-D were significantly lower in DHEA+ relative to DHEA. Collectively, our data suggest a role of DHEA on these metabolic and cytokines pathways, and these FF metabolites could be used to guide future studies in DHEA supplementation regimen.

## Introduction

Poor ovarian responders (POR) are a sub-group of infertile women that account for 9-26% of *in vitro* fertilization (IVF) indications.^1,2^ In patients designated as “poor responders,” so-called due to poor response to ovarian stimulation given during IVF workup, the limited number of obtained oocytes remains the major problem in optimizing the live birth rates.^3^ In fact, as a result of a lower number of oocytes retrieved, there are fewer embryos to select and transfer, and subsequently these patients have lower pregnancy rates per transfer and lower cumulative pregnancy rates per started cycle compared with normal responders. In PORs the mechanism of ovarian insufficiency can be multifactorial with causes such as ovarian surgery especially in case of endometrioma,^4,5^ uterine artery embolization for the treatment of uterine leiomyoma,^6,7^ genetic defects, chemotherapy, radiotherapy, autoimmune disorders, single ovary, chronic smoking,^8,9^ or linked to diseases such as diabetes mellitus Type I.^10^ However, in most cases, follicular depletion plausibly reflecting premature ovarian aging,^11^ which clinically translates into a reduction of implantation rates, an increase of early pregnancy loss, and disappointingly low IVF success.^12,13^ In each menstrual cycle, human ovaries produce a single dominant follicle. Growth of the dominant follicle encompasses enlargement of the oocyte, replication of follicular cells, and formation and expansion of a fluid-filled follicular antrum or cavity, providing a specialized microenvironment for the development of oocytes. Follicular fluid (FF) that fills the antrum cavity is derived from the surrounding theca capillaries, abundant and easily accessible during IVF procedures due to ample volume being produced during follicle maturation.^14^ FF are rich in metabolites, notably hormones, and proteins that are critical for oocyte growth and development, which determines subsequent potential to achieve fertilization and embryo development. As such, constituents of the FF surrounding the oocyte provides a unique biochemical window about the growth and differentiation of the oocyte.^15^ To divulge into the biochemical space of human FF, gas chromatography-mass spectrometric (GC-MS) and proton nuclear magnetic resonance (^1^H NMR) metabolomic analyses have been previously conducted,^16–19^ as were proteomic analyses.^20–25^ These studies mainly report the IVF FF profile, whereas the FF metabolome of POR remains poorly characterized. Dehydroepiandrosterone (DHEA) is a steroid produced in the adrenal cortex and the ovarian theca cells in women that is converted into more active forms of androgens such as testosterone.^5^ It has been suggested that DHEA supplementation may increase the number of available follicles in PORs through an increased serum level of insulin-like growth factor, increased follicular response to follicle stimulating hormone (FSH), and improved quality of oocytes.^26^ However, the efficacy of DHEA pre-treatment has been controversial, with partial to low clinical evidence being observed.^9,13–14,16^ Here, a randomized clinical trial was conducted to evaluate the effects of DHEA on IVF outcomes in our patient population, and a metabolomics approach was used to understand the effects of DHEA on the FF metabolome. Given that exogenous stimulus such as DHEA supplementation might alter the FF metabolic profile, we hypothesized that DHEA effects could be improved through uncovering ‘responder’ FF metabolites, which could be tapped to improve DHEA dose regime. With the advent of LC-MS/MS, LC-MS global (untargeted) metabolomics analysis has provided the ability to reveal biologically relevant changes within a system, even at sensitive ranges before the precedence of gross morphological or phenotypical changes.^28^ Furthermore, DHEA has immunoregulatory functions, and the large-scale study of cytokines plausibly reveals DHEA immunomodulatory targets.^29^ A better understanding of the molecular mechanisms implicated in the conversion of DHEA to testosterone will offer investigators the opportunity to design individualized treatments for infertile women with POR, which are tailored to patients’ unique metabolic profiles. Therefore, in this study we mapped the FF metabolome and cytokine profile of POR with and without DHEA supplementation.

## Material and methods

### Ethical approval

The study was approved by the Institutional Review Board at the Singapore Health Systems, and written informed consent was obtained from each participant. The study was registered with the National Institutes of Health (NIH) clinical trial site (clinicaltrials.gov: NCT01535872).

### Study population and design

The prospective open-label randomized controlled trial (RCT) was conducted to evaluate the effect of DHEA administration in women, below the age of 42 starting their IVF treatment who met one of the two following features of POR (an abnormal ovarian reserve test and/or a previous poor response to ovarian stimulation in an IVF cycle) were assessed for eligibility.^3^ We defined POR in this study as women with diminished ovarian reserves (AMH <1.0 ng/mL or Day 2 or 3 FSH >10 IU/L), or women with fewer than four oocytes retrieved with either standard long or antagonist protocols.^3^ Inclusion criteria included women with diminished ovarian reserves (anti-müllerian hormone <1.0 ng/mL or D2/3 follicle stimulating hormone >10 IU/L), or women with fewer than four oocytes retrieved with either standard long or antagonist protocols. Exclusion criteria included women with previous or current DHEA supplementation, use of corticosteroids within the past three months, major systemic illnesses, and allergy to DHEA. Eligible patients were randomized to either the DHEA treatment group (DHEA+, *N*=28) and received DHEA (Pharma Natural, USA) at the dose of 75 mg/day for three to eight months prior starting their controlled ovarian stimulation (COS), or the control group, receiving no treatment (DHEA-, *N*=24) (**Table 1**). Average age (DHEA-, 36 years; DHEA+, 37 years) and body mass index (DHEA-, 24 kg/m^2^; DHEA+, 23 kg/m^2^) were similar in both groups (*p*>0.05).

**Table 1.** Baseline characteristics of patients in this study

|  | **DHEA group (n = 28)** | **Control group (n = 24)** |
| --- | --- | --- |
| Age (year), mean (SD) | 37 (3) | 36 (4) |
| BMI (kg/m<sup>2</sup>), mean (SD) | 23 (5) | 24 (5) |
| Race, n (%) |  |  |
| Chinese | 21 (75.0) | 14 (58.3) |
| Malay | 0 (0.0) | 8 (33.3) |
| Indian | 1 (3.6) | 2 (8.4) |
| Others | 6 (21.4) | 0 (0.0) |
| Primary infertility, n (%) |  |  |
| Yes | 19 (67.9) | 13 (54.2) |
| No | 9 (32.1) | 11 (45.8) |
| Infertility duration (years), median (range) | 4 (1-16) | 4 (1-11) |
| Primary infertility diagnosis, n (%) |  |  |
| Male factor | 22 (78.6) | 17 (70.8) |
| Tubal factor | 1 (3.6) | 1 (4.2) |
| Endometriosis | 1 (3.6) | 2 (8.3) |
| Low ovarian reserve | 3 (10.6) | 4 (16.7) |
| Others | 1 (3.6) | 0 (0.0) |
| Cycle number n (%) |  |  |
| Cycle 1 | 4 (14.3) | 12 (50.0) |
| Cycle 2 & above | 22 (78.6) | 12 (50.0) |
| Basal FSH (IU/L) , mean (range)* | 8.4 (4.8-27.7) | 6.5 (3.6-16.6) |
| Estradiol (pmol/L) , mean (range)* | 98.7 (37.0-320.0) | 92.4 (37.0-273.0) |
| AMH (ng/ml) , mean (range)* | 0.7 (0.2-2.8) | 0.8 (0.2-2.7) |
| Free Testosterone (pmol/L), mean (range)* | 2.0 (0.5-14.3) | 1.7 (0.9-2.9) |
| DHEAs (μmol/L), mean (range)* | 4.0 (0.5-17.3) | 3.8 (0.9-10.0) |
| Antral follicle count, mean (range)* | 4.6 (1-15) | 5.1 (0-12) |
| Ovarian volume RO (cm<sup>3</sup>), mean (range)* | 7.3 (1.9-24.0) | 7.7 (2.0-29.5) |
| Ovarian volume LO (cm<sup>3</sup>), mean (range)* | 5.1 (1.8-16.6) | 8.2 (2.4-26.7) |

\* Blood baseline parameters, antral follicle count and ovarian volumes were obtained in 27 patients in the DHEA group and 20 patients in the control group and were summarized by geometric mean (range).

### IVF/ICSI protocol

All individuals received the same stimulation protocol, same starting dose of gonadotropin, and fertilization technique. Briefly, the IVF/ICSI treatment cycle involved an antagonist based COS protocol consisting of daily sub-cutaneous injections of recombinant-FSH (Puregon, Follitropin beta, 300iu; MSD, USA) and highly-purified human menopausal gonadotropin (Menopur; Menotropin, 150 IU; Ferring Pharmaceuticals, Germany) with initiation of gonadotropin releasing hormone antagonist (Ganirelix, Orgalutan, 0.25 mg s/c; MSD, USA) on day 5 of COS. The dose of Menopur and Puregon could be further increased depending on individual ovarian response. All patients had this standardized antagonist (short) protocol: no agonist (long) protocol was used. Human chorionic gonadotropin (i.m 10,000 IU hCG; Pregnyl; MSD, USA) was administered when at least one follicle measured ≥ 17 mm in diameter (averaged orthogonal measurements). The endometrial thickness, peak estradiol and progesterone levels were assessed on the day of human chorionic gonadotropin (hCG) trigger. Ultrasound-guided trans-vaginal oocyte retrieval was performed 36 hours after hCG administration. The effect of DHEA supplementation on the markers of ovarian reserve (anti-müllerian hormone; AMH), and follicular function (IGF-1)^30^, as well as ovarian follicular levels of estradiol, testosterone, and DHEA collected from the lead follicle at the time of OPU were assessed through ELISA as previously described.^31^

The embryo transfer was performed on day 2 or day 3 of embryo-culture, and luteal phase support was achieved with vaginal progesterone (micronized progesterone, Utrogestan, 200 mg three times a day, Besins-International, France). Pregnancy was established by serum beta-hCG seventeen days post embryo transfer. Clinical pregnancy will be established by a transvaginal ultrasound four weeks after embryo transfer. IVF/ICSI primary and secondary outcomes are shown in **Table 3**.

### Sample preparation

For untargeted metabolomics analysis, sample preparation followed a previously published report with some modifications.^32^ FF were available for metabolomics and cytokine analyses (DHEA+, *N*=18 and DHEA-, *N*=16). A volume of 50 μL from each FF sample was thawed at 4°C, and FF proteins were precipitated with 200 μL ice-cold methanol. After vortexing, the mixture was centrifuged at 16,000 rpm for 10 min at 4°C and the supernatant was collected and evaporated to dryness in a speedvacuum evaporator. The dry extracts were then re-dissolved in 200 μL of water/methanol (98:2; v/v) for liquid chromatography-tandem mass spectrometry (LC-MS/MS) analysis.

A pooled quality control (QC) sample was generated to allow comparison of analytic behavior over long periods of time. The pooled reference samples were for the purposes of quality control (i.e., to ensure relative consistency among identical samples within days) and for quality assurance (i.e., to ensure consistent results between days). They did not contribute data to downstream statistical analysis.

### Liquid Chromatography-Tandem Mass Spectrometry-based Metabolomics

The supernatant fraction from sample preparation step was analyzed using Agilent 1290 ultra-high pressure (performance) liquid chromatography system (Waldbronn, Germany) equipped with a Bruker impact II Q-TOF mass spectrometer with its normal electrospray ionization (ESI) ion source (Bruker Daltonics). 2.5 μL of samples was injected and were separated using Waters Acquity HSS T3 (2.1 mm i.d. x 100 mm, 1.8 μm) at a flow rate of 0.2 mL/min. The oven temperature was set at 50°C. The gradient elution involved a mobile phase consisting of (A) 0.1% formic acid in water and (B) 0.1% formic acid in methanol. The initial condition was set at 5% B. A 5.5 min linear gradient to 60% B was applied, followed by a 13 min gradient to 98% B (total 24 min including wash and re-equilibration) at a flow rate of 0.4 ml/min. The ion spray voltage was set at 4,500 V, and the Dry Temperature was maintained at °C. The drying nitrogen gas flow rate and the nebulizer gas pressure were set at 8.0 L/min and 26 psi, respectively. Calibration of the system was performed using sodium formate clusters before data acquisition. The stability of the LC-MS method was examined and evaluated by sodium formate clusters (1 mM NaOH, 0.1% formic acid, 50% 2-propanol) infused into the system.

The ESI mass spectra were acquired in positive ion mode. Mass data were collected between *m/z* 100 and 1000 at a rate of three scans per second. Auto MS/MS was triggered at 8 Hz with duty cycle of 1.5 s. Threshold was set at 1500 counts, with active exclusion activated after 3 spectra, released after 0.3 min and overwritten if the current or previous intensity changes. MS/MS spectra were acquired at collision energy of 20–50 eV automatically varied by the charge states and the intensities of the selected precursors. Fragment spectra acquisition was carried out at a scan rate dependent on the MS precursor intensities - MS/MS spectra for high-intensity precursors were acquired for a shorter time (90000 counts, 12 Hz) than low-intensity precursor ions (10000 counts, 6 Hz) thus allowing for a balancing of maximal scan time and MS/MS spectral quality.

### Compound identification

Structure identification was achieved via the following in MetaboScape (version 2.0; **Figure 1**): elemental composition was predicted via isotopic pattern following the rules (i) mSigma of MS1: 20 with tolerance of 5 ppm and (ii) MS2: 50 with tolerance of 2 mDa of the differential metabolites was searched against Bruker HMDB (Human Metabolome Database) using a precursor match of ±10 mDa, minimum score of 400 and minimum match score of 250. Progesterone, glycerophosphocholinelinoleic acid and valine were structurally confirmed using chemical standards.

**Figure 1.**
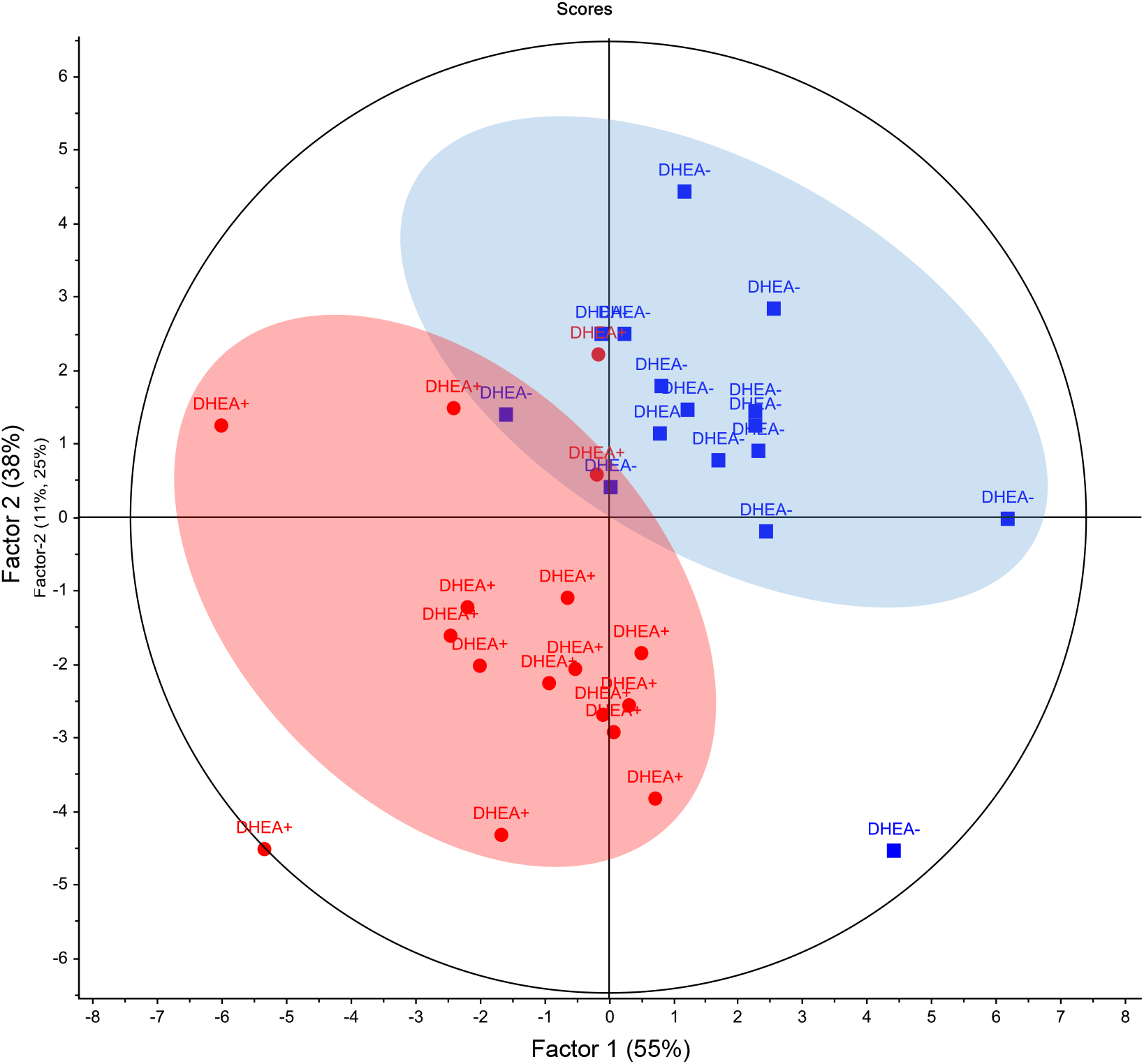
Partial Least Squares Scores plot of DHEA- and DHEA+ follicular fluid metabolome. Metabolomic data was median centred, and scaled by division with the standard deviation. The follicular fluid metabolome distinguished poor ovarian response subjects on DHEA supplementation (DHEA+, red) and without DHEA supplementation (DHEA-, blue).

### Multiplex immunoassay analysis

45 cytokines were detected and measured using ProCartaplex (EBioscience, CA, USA) as previously reported [BDNF, EGF, Eotaxin (CCL11), FGF-2 (FGF basic), GM-CSF, CXCL1 (GROα), HGF, IFNγ, IFNα, IL-1RA, IL-1β, IL-1α, IL-2, IL-4, IL-5, IL-6, IL-7, CXCL8 (IL-8), IL-9, IL-10, IL-12 p70, IL-13, IL-15, IL-17A, IL-18, IL-21, IL-22, IL-23, IL-27, IL-31, CXCL10 (IP-10), LIF, CCL2 (MCP-1), CCL3 (MIP-1α), CCL4 (MIP-1β), βNGF, PDGF-BB, PLGF, CCL5 (RANTES), SCF, CXCL12 (SDF1α), TNFα, LTA (TNFβ), VEGF-A, VEGF-D].^33^ Briefly, 5 μL of FFs were diluted with 5 μL Universal Dilution Buffer, and mixed with 50 μL of antibody-conjugated, magnetic beads in a 96 well DropArray plate (Curiox Biosystems, Singapore) and rotated at 450 rpm for 120 min at 25°C while protected from light. Beads were internally dyed with different concentrations of two spectrally distinct fluorophores and covalently conjugated to antibodies against the 45 cytokines, chemokines and growth factors. The plate was washed three times with wash buffer (PBS, 0.05% Tween-20) on the LT210 Washing Station (Curiox) before adding 10 μL of secondary antibody and rotating at 450 rpm for 30 min at 25°C protected from light. Subsequently, the plate was washed three times with wash buffer, and 10 μL of streptavidin-phycoerythrin added and rotated at 450 rpm for 30 min at 25°C protected from light. The plate was again washed thrice with wash buffer; 60 μL of reading buffer was then added and the samples read using the Bio-Plex Luminex 200 (BioRad). The beads were classified by the red classification laser (635 nm) into its distinct sets, while a green reporter laser (532 nm) excites the phycoerythrin, a fluorescent reporter tag bound to the detection antibody. Quantitation of the 45 cytokines was then determined by extrapolation to a six or seven-point standard curve using five-parameter logistic regression modelling. Calibrations and validations were performed prior to runs and on a monthly basis respectively.

### Statistical analysis

GraphPad Prism 6 (GraphPad Software Inc.) was used for performing all statistical analyses. Data were checked for normal distribution using Kolmogorov-Smirnov test. Unpaired or paired t-test was performed, as appropriate, to determine statistical significance between groups form normally distributed data. Mann-Whitney U test was used for non-normally distributed data. For comparing more than three groups, the data were analyzed using ANOVA test, followed by the *t*-test with Bonferroni adjustment. *P*<0.05 was considered significant. Metabolomic data was further analyzed by Principal Component Analysis (PCA) and Partial Least Squares Regression (PLSR) modelling (Unscrambler X version 10.1) after the normalization of data by first centering the data to the median and scaling it by division with the standard deviation. Full cross-validation was applied in PLSR to increase model performance and for the calculation of coefficient regression values.

## Results

### LC-MS/MS metabolomics system evaluation

In order to obtain reliable metabolic profiles of the samples, we evaluated the stability and reproducibility of the LC-MS method by performing Principal Component Analysis (PCA) on all the samples including the eight pooled quality control samples. As shown in **Figure S1**, the QC samples clustered in PCA scores plots, and together with retention time CV% < 0.1 min, peak *m/z* values 3 mDa, and relative standard deviations of peak areas < 20% (**Table 1**), there was good system stability, mass accuracy and reproducibility of the chromatographic separation during the whole LC-MS/MS sequence. In addition, intensity CV% of the identified compounds in pooled quality control samples are low (average 6%; **Table 1**). PCA hotelling (T^2^) revealed one DHEA+ subject as an outlier (D4) and was removed from further analysis (**Figure S1**). From a total 2717 time-aligned features, an average of 903 features was chosen for auto MS/MS mode. From these, a total of 100 metabolites were identified via chemical standard confirmed HMDB.^34^ An average of 65 MS/MS confirmed metabolites per patient was identified, which was similar in terms of metabolite identified in either DHEA- or DHEA+ subjects (*p*=0.8; range:59-76 metabolites; **Table S1**).

### POR follicular fluid metabolome

In POR subjects, the FF metabolome spanned three orders of magnitude, and was composed of a diversity of metabolites including, glycerophospholipids and derivatives (glycerophosphocholine, phosphatidylcholines), fatty acids (heptadecanoic acid, linoleic acid, vaccenic acid, myristic acid), cholesterols (isocaproic acid, 7-ketocholesterol), glucocorticoids (11-deoxycortisol or cortexolone, cortisol, corticosterone), hormones (17-hydroxyprogesterone, deoxycorticosterone, 11α-hydroxyprogesterone, 16-dehydroprogesterone, androstenedione, epitestosterone, progesterone, pregnenolone). Other metabolites included bile acids (3b-hydroxy-5-cholenoic acid, 3-oxocholic acid, glycocholic acid), peptides and derivatives (3-indolepropionic acid), lactones (delta-hexanolactone/caprolactone), lactic acid, vitamin D3 and sphingosine (**Table 2**).

**Table 2.** List of identified follicular fluid metabolites

| No. | HMDB No. | Accurate mass | Theoretical mass | Compound Name | Chemical formula | Pathway |
| --- | --- | --- | --- | --- | --- | --- |
| 1 | HMDB01859 | 151.0626 | 151.0633 | Acetaminophen | C<sub>8</sub>H<sub>9</sub>NO<sub>2</sub> | Acetaminophen metabolism |
| 2 | HMDB04987 | 261.1206 | 261.1325 | Alpha-Aspartyl-lysine | C<sub>10</sub>H<sub>19</sub>N<sub>3</sub>O<sub>5</sub> | Amino acid metabolism |
| 3 | HMDB03423<br>/HMDB00641 | 146.0687 | 146.0691 | D-Glutamine/L-Glutamine | C<sub>5</sub>H<sub>10</sub>N<sub>2</sub>O<sub>3</sub> | Amino acid metabolism |
| 4 | HMDB00714 | 179.0576 | 179.0582 | Hippuric acid | C<sub>9</sub>H<sub>9</sub>NO<sub>3</sub> | Amino acid metabolism |
| 5 | HMDB00161<br>/HMDB01310/<br>HMDB00271 | 89.0465 | 89.0477 | L-Alanine/D-Alanine/Sarcosine | C<sub>3</sub>H<sub>7</sub>NO<sub>2</sub> | Amino acid metabolism |
| 6 | HMDB00641<br>/HMDB03423 | 146.0687 | 146.0691 | L-Glutamine/ D-Glutamine | C<sub>5</sub>H<sub>10</sub>N<sub>2</sub>O<sub>3</sub> | Amino acid metabolism |
| 7 | HMDB00687<br>/HMDB00557/<br>HMDB00172/<br>HMDB01645 | 131.0941 | 131.0946 | L-Leucine/L-Alloisoleucine/ L-Isoleucine/L-Norleucine | C<sub>6</sub>H<sub>13</sub>NO<sub>2</sub> | Amino acid metabolism |
| 8 | HMDB00696 | 149.0504 | 149.0510 | L-Methionine | C<sub>5</sub>H<sub>11</sub>NO<sub>2</sub>S | Amino acid metabolism |
| 9 | HMDB00162 | 115.0629 | 115.0633 | L-Proline | C<sub>5</sub>H<sub>9</sub>NO<sub>2</sub> | Amino acid metabolism |
| 10 | HMDB00883<br>/HMDB00043 | 117.0784 | 117.0790 | L-Valine/Betaine | C<sub>5</sub>H<sub>11</sub>NO<sub>2</sub> | Amino acid metabolism |
| 11 | HMDB00064<br>/HMDB00766 | 131.0688 | 131.0695 | Creatine/ N-Acetyl-L-alanine | C<sub>4</sub>H<sub>9</sub>N<sub>3</sub>O<sub>2</sub> | Arginine, proline, glycine and serine metabolism |
| 12 | HMDB00043<br>/HMDB02141 | 117.0786 | 117.0790 | Betaine/ N-Methyl-a-aminoisobutyric acid | C<sub>5</sub>H<sub>11</sub>NO<sub>2</sub> | Betaine Metabolism |
| 13 | HMDB01847 | 194.0796 | 194.0804 | Caffeine | C8H10N4O2 | Caffeine metabolism |
| 14 | HMDB01860 | 180.0642 | 180.0647 | Paraxanthine | C7H8N4O2 | Caffeine metabolism |
| 15 | HMDB02825 | 180.0638 | 180.0647 | Theobromine | C7H8N4O2 | Caffeine metabolism |
| 16 | HMDB00062 | 161.1048 | 161.1052 | L-Carnitine | C7H15NO3 | Fatty acid metabolism |
| 17 | HMDB00267<br>/HMDB0080<br>5 | 129.0419 | 129.0426 | Pyroglutamic acid/<br>Pyrrolidonecarboxylic acid | C5H7NO3 | Glutathione metabolism |
| 18 | HMDB00017 | 183.0528 | 183.0532 | 4-Pyridoxic acid | C8H9NO4 | Vitamin B6 metabolism |
| 19 | HMDB00995 | 312.2087 | 312.2089 | 16-Dehydroprogesterone | C21H28O2 | Lipid metabolism |
| 20 | HMDB00502 | 388.2485 | 406.2719 | 3-Oxocholic acid | C24H38O5 | Lipid metabolism |
| 21 | HMDB00308 | 356.2708 | 374.2821 | 3b-Hydroxy-5-cholenoic acid | C5H4N4O3 | Lipid metabolism |
| 22 | HMDB00501 | 400.3339 | 400.3341 | 7-Ketocholesterol | C27H44O2 | Lipid metabolism |
| 23 | HMDB00503 | 372.2659 | 390.2770 | 7a-Hydroxy-3-oxo-5b-<br>cholanoic acid | C24H38O4 | Lipid metabolism |
| 24 | HMDB00784 | 188.1041 | 188.1049 | Azelaic acid | C9H16O4 | Lipid metabolism |
| 25 | HMDB00015<br>/HMDB0154<br>7 | 346.2131 | 346.2144 | Cortexolone/Corticosterone | C21H30O4 | Lipid metabolism |
| 26 | HMDB01547 | 346.2142 | 346.2144 | Corticosterone | C21H30O4 | Lipid metabolism |
| 27 | HMDB00063 | 362.2093 | 362.2093 | Cortisol | C21H30O5 | Lipid metabolism |
| 28 | HMDB00631 | 449.3131 | 449.3141 | Deoxycholic acid glycine<br>conjugate | C26H43NO5 | Lipid metabolism |
| 29 | HMDB00573<br>/HMDB0323<br>1 | 282.2564 | 282.2559 | Elaidic acid/Vaccenic acid | C18H34O2 | Lipid metabolism |
| 30 | HMDB00628<br>/HMDB0023<br>4 | 288.2078 | 288.2089 | Epitestosterone/Testosteron<br>e | C19H28O2 | Lipid metabolism |
| 31 | HMDB00086 | 257.1026 | 257.1028 | Glycerophosphocholine | C8H20NO6P | Lipid metabolism |
| 32 | HMDB00138 | 465.3096 | 465.3090 | Glycocholic acid | C26H43NO6 | Lipid metabolism |
| 33 | HMDB02259 | 270.2557 | 270.2559 | Heptadecanoic acid | C17H34O2 | Lipid metabolism |
| 34 | HMDB00666 | 130.0989 | 130.0994 | Heptanoic acid | C7H14O2 | Lipid metabolism |
| 35 | HMDB00689 | 116.0833 | 116.0837 | Isocaproic acid | C6H12O2 | Lipid metabolism |
| 36 | HMDB00673 | 280.2397 | 280.2402 | Linoleic acid | C18H32O2 | Lipid metabolism |
| 37 | HMDB00806 | 228.2088 | 228.2089 | Myristic acid | C14H28O2 | Lipid metabolism |
| 38 | HMDB00593 | 785.5943 | 785.5935 | PC(18:1/18:1) | C44H84NO8P | Lipid metabolism |
| 39 | HMDB00847 | 158.1300 | 158.1307 | Pelargonic acid | C9H18O2 | Lipid metabolism |
| 40 | HMDB00253 | 316.2395 | 316.2402 | Pregnenolone | C21H32O2 | Lipid metabolism |
| 41 | HMDB01830 | 314.2239 | 314.2246 | Progesterone | C21H30O2 | Lipid metabolism |
| 42 | HMDB00792 | 202.1194 | 202.1205 | Sebacic acid | C10H18O4 | Lipid metabolism |
| 43 | HMDB00933 | 228.1335 | 228.1362 | Traumatic acid | C12H20O4 | Lipid metabolism |
| 44 | HMDB03231 | 282.2551 | 282.2559 | Vaccenic acid | C18H34O2 | Lipid metabolism |
| 45 | HMDB01877 | 144.1146 | 144.1150 | Valproic acid | C8H16O2 | Lipid metabolism |
| 46 | HMDB00876 | 384.3387 | 384.3392 | Vitamin D3 | C27H44O | Lipid metabolism |
| 47 | HMDB00182<br>/HMDB0340<br>5 | 146.1048 | 146.1055 | L-lysine/D-lysine | C6H14N2O2 | Lysinuric protein intolerance |
| 48 | HMDB01923 | 230.0937 | 230.0943 | Naproxen | C14H14O3 | Naproxen action pathway |
| 49 | HMDB06344 | 264.1103 | 264.1110 | Alpha-N-phenylacetyl-L-glutamine | C13H16N2O4 | Phenylacetate Metabolism |
| 50 | HMDB00159 | 165.0784 | 165.1891 | L-Phenylalanine | C9H11NO2 | Phenylalanine metabolism |
| 51 | HMDB00097 | 103.0994 | 104.1075 | Choline | C5H14NO | Phosphatidylcholine biosynthesis |
| 52 | HMDB00157 | 136.0380 | 136.0385 | Hypoxanthine | C5H4N4O | Purine metabolism |
| 53 | HMDB00289 | 168.0278 | 168.0283 | Uric acid | C5H4N4O3 | Purine metabolism |
| 54 | HMDB00926 | 79.0421 | 79.0422 | Pyridine | C5H5N | Pyridine biosynthesis |
| 55 | HMDB00975<br>/HMDB00055 | 324.1052 | 342.1162 | Trehalose/Cellobiose | C12H22O11 | Pyrimidine metabolism |
| 56 | HMDB00300 | 112.0266 | 112.0273 | Uracil | C4H4N2O2 | Pyrimidine metabolism |
| 57 | HMDB00190<br>/HMDB01311 | 90.0311 | 90.0317 | L-Lactic acid/D-Lactic acid | C3H6O3 | Pyruvate metabolism |
| 58 | HMDB00252 | 299.2818 | 299.2824 | Sphingosine | C18H37NO2 | Sphingolipid Metabolism |
| 59 | HMDB00374<br>/HMDB00016/HMDB00920 | 330.2178 | 330.2195 | 17-Hydroxyprogesterone/Deoxycorticosterone/11a-Hydroxyprogesterone | C21H30O3 | Steroid biosynthesis |
| 60 | HMDB00929 | 204.0894 | 204.0899 | L-Tryptophan | C11H12N2O2 | Tryptophan metabolism |
| 61 | HMDB00197 | 175.0627 | 175.0633 | Indoleacetic acid | C10H9NO2 | Tryptophan metabolism |
| 62 | HMDB00183 | 208.0843 | 208.0848 | L-Kynurenine | C10H12N2O3 | Tryptophan metabolism |
| 63 | HMDB00158<br>/HMDB06050 | 181.0735 | 181.0739 | L-Tyrosine/o-Tyrosine | C9H11NO3 | Tyrosine metabolism |
| 64 | HMDB02302 | 189.0782 | 189.0790 | 3-Indolepropionic acid | HMDB02302 | Tryptophan deamination |
| 65 | HMDB01924 | 266.1620 | 266.1630 | Atenolol | C14H22N2O3 | Beta1-receptor inhibition |
| 66 | HMDB06115 | 106.0416 | 106.0419 | Benzaldehyde | C7H6O | Oxidoreductase activity |
| 67 | HMDB00562 | 113.0586 | 113.0589 | Creatinine | C4H7N3O | Arginine, proline, glycine and serine Metabolism/creatine catabolism |
| 68 | HMDB00453 | 114.0670 | 114.0681 | Delta-hexanolactone | C6H10O2 | Hydroxy acid lactonization |
| 69 | HMDB04983 | 94.0084 | 94.0089 | Dimethyl sulfone | C2H6O2S | Methanethiol metabolism |
| 70 | HMDB01888 | 73.0524 | 73.0528 | N,N-Dimethylformamide | C3H7NO | Tertiary carboxylic acid metabolism |
| 71 | HMDB00070 | 129.0785 | 129.0790 | Pipecolic acid | C6H11NO2 | Amino acid metabolism |

**Table 3.**
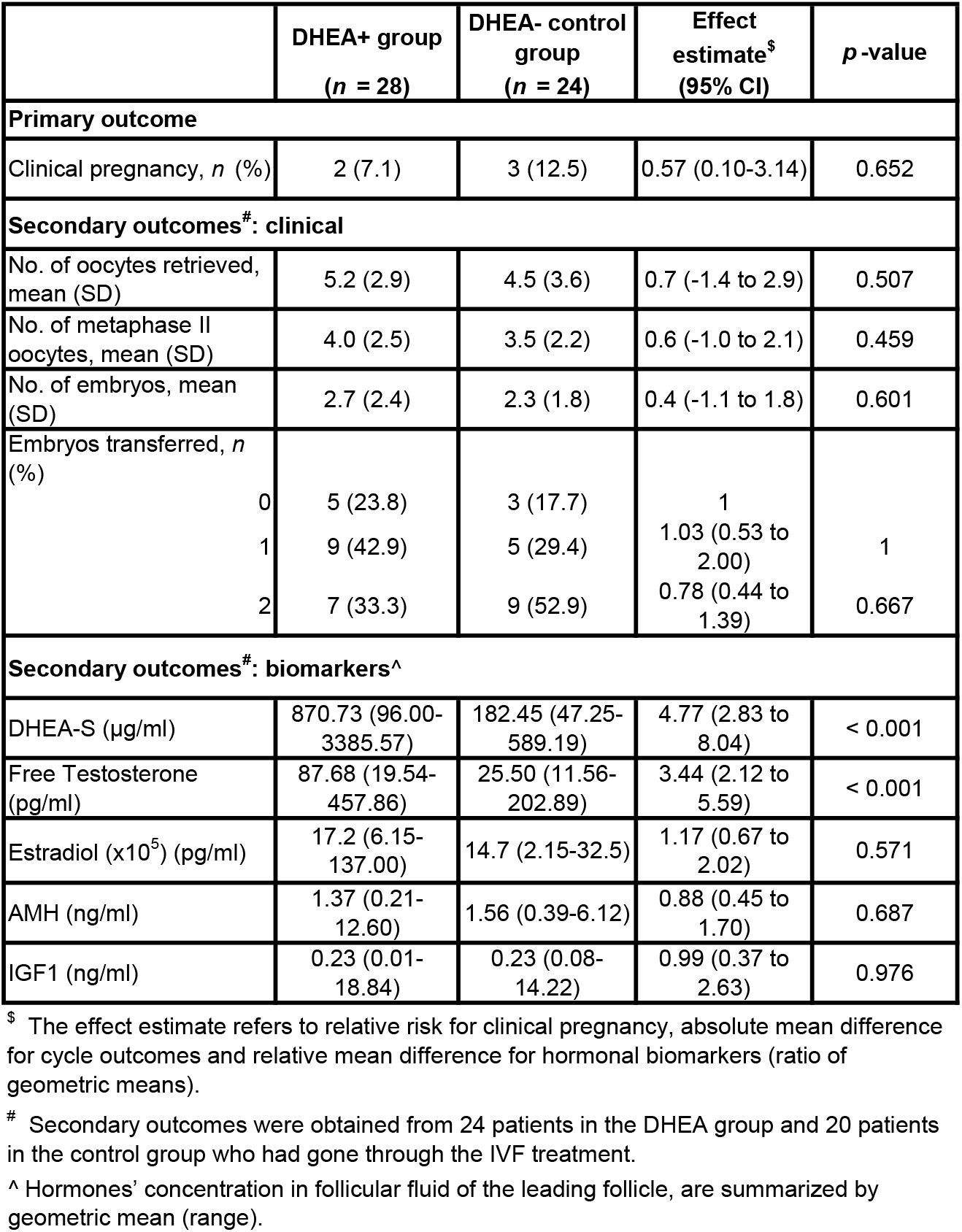
Primary and secondary outcomes between DHEA+ and DHEA- control groups.

### Altered follicular fluid metabolome in response to DHEA

Between the DHEA+ and DHEA- groups, the FF metabolome profiles were distinct. Based on the FF metabolomes, PLSR modelling distinguished the DHEA+ subjects from the controls at 2 PLS components, describing 55% of the X-variance and 38% of the Y-variance (**Figure 1**). In DHEA- subjects, progesterone was the most abundant FF metabolite, followed by L-alanine, L-phenylalanine, pyridine, L-leucine, which the top five metabolites collectively made up close to half (48%) of the FF metabolome (**Figure 2A**). In DHEA+ subjects, the FF metabolome profile of highly abundant metabolites was different, with cortisol as the most abundance metabolite, followed by L-alanine, L-phenylalanine, pyridine, L-isoleucine and L-leucine. These top six metabolites collectively made up ~49.5% of the DHEA+ FF metabolome (**Figure 2B**). Interestingly, pyridine considered a non-endogenous metabolite (HMDB0000926), was found in such abundance which suggested it came from the synthesis of DHEA.^35^ Notably, the observed MS/MS spectra of pyridine at various eV matched very well with HMDB database (**Figure S2A**), which suggested its correct identification. As a precursor to testosterone and estrogen, DHEA could be converted to testosterone, and aromatized to estrogen; in the case of POR, exogenous DHEA was proposed to increase androgens in promoting folliculogenesis and potentiate the effects of gonadotropins.^8,36,37^ FF testosterone was detected in our metabolomics profiling, although the differences between DHEA+ and DHEA- group were small [DHEA-: mean signal intensity= 2294.5; DHEA+: mean signal intensity=2267.75 (testosterone), *p*>0.05; **Figure S2B**].

**Figure 2.**
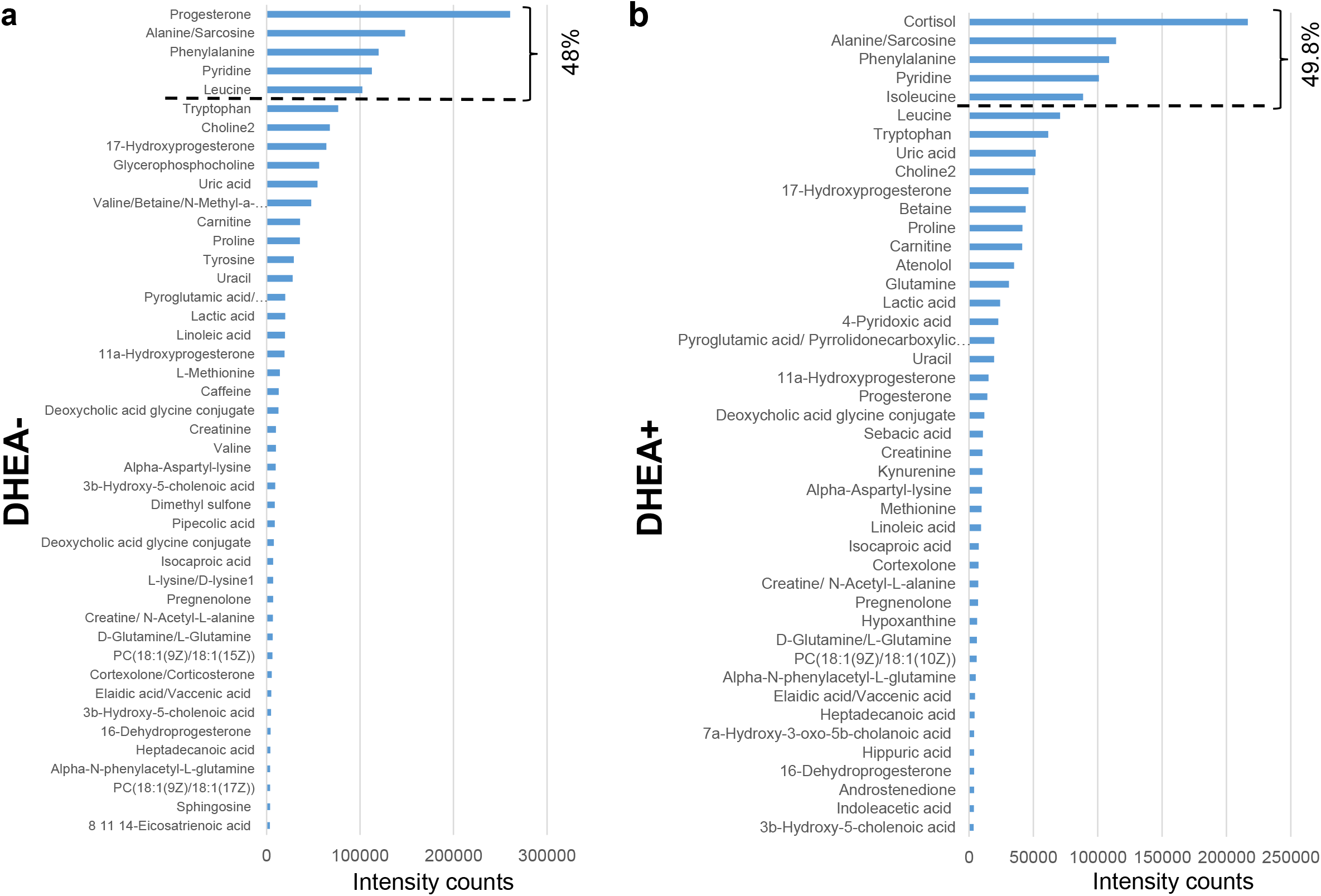
Histogram of follicular fluid metabolites in poor ovarian responders with and without DHEA supplementation. Follicular metabolome coverage and metabolite abundance as quantified by untargeted LC-MS/MS metabolomics in (a) DHEA- and (b) DHEA+ subjects, and were ranked according to their intensity counts. The top 50% metabolites are shown, with progesterone being the major differentiating metabolite between the DHEA- and DHEA+ subjects.

Four FF metabolites, namely, glycerophosphocholine, linoleic acid, progesterone, and L-valine were significantly lower in DHEA+ relative to DHEA- (*p*<0.05-0.005; **Figure 3A-D**). Interestingly, pregnenolone, a cholesterol metabolite and steroid that is upstream of DHEA metabolism, was detected only in DHEA+ (6/18 subjects), and not DHEA- (0/16 subjects). Receiver operating characteristic (ROC) analyses of the four metabolites revealed area under the curve (AUC) ranging from 0.711 (progesterone), 0.730 (glycerophosphocholine), 0.785 (linoleic acid) and 0.818 (L-valine) (*p*<0.05-0.01; **Figure 3E-H**), suggesting the utility of these FF metabolites in monitoring DHEA dose modulation. Linoleic acid and L-valine remained significantly lower in DHEA+ (*p*<0.05, *p*<0.001 for both) when women with endometriosis (*N*=5) were removed from analysis, strongly suggesting the significant effect of DHEA on these metabolites (**Figure S3**).

**Figure 3.**
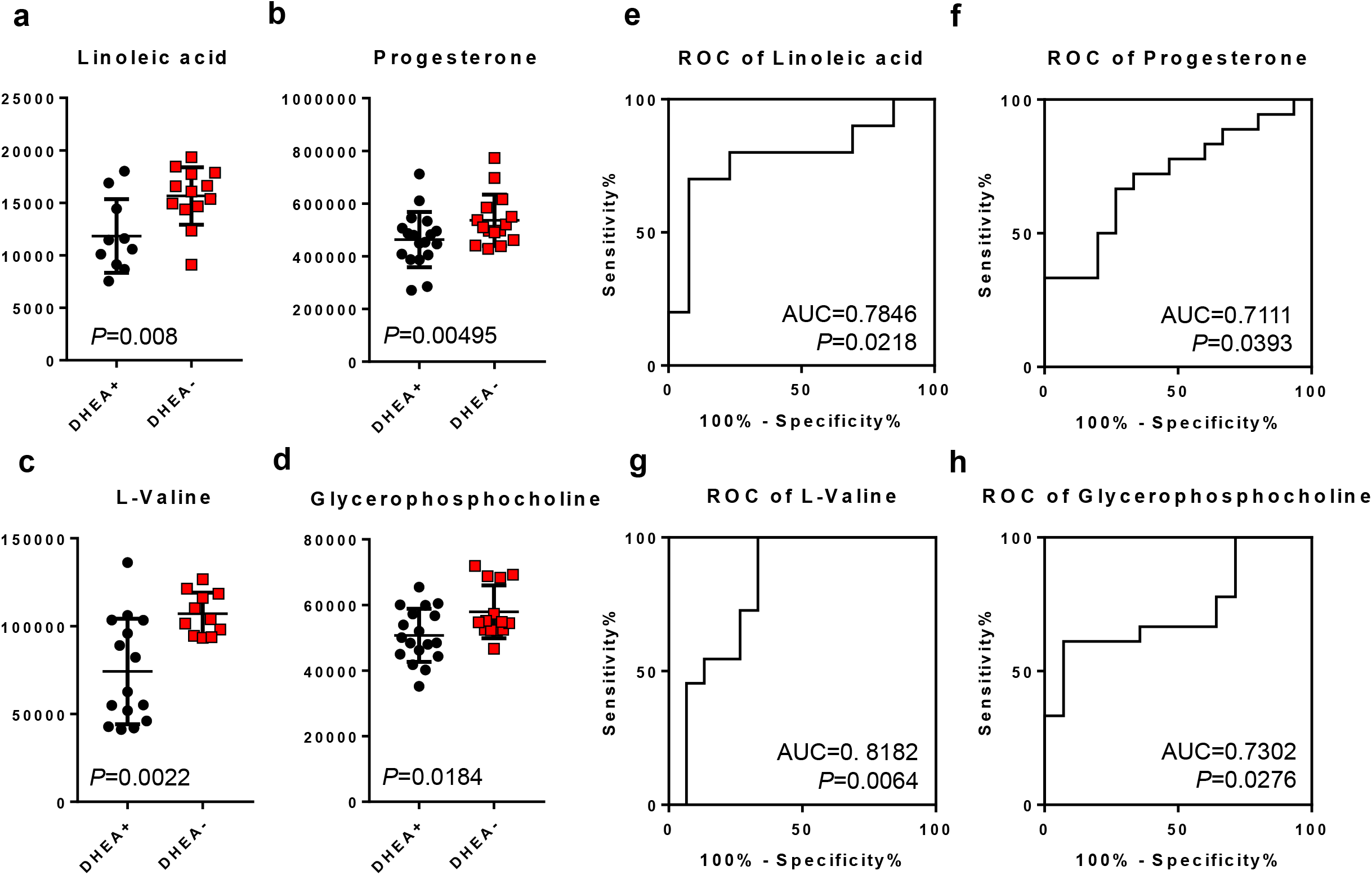
Significantly different follicular fluid metabolites in DHEA+ and DHEA-patients. (A-D) Dot plots of significantly different FF metabolites. Student’s t-tests were performed and p<0.05 is considered statistically significant. (E-F) corresponding receiver operating curve (ROC) analyses of the metabolites. Area under curve (AUC) of the metabolites and their P-values are reported.

### FF cytokine profile in response to DHEA

Among the 45 immunomodulatory proteins (cytokines, chemokines and growth factors), 22 were detected in human FF, namely 10 cytokines (IFNγ, IL12p70, IL13, IL1b, TNFα, IL1Ra, IL5, IL7, IL10, IL18), 6 chemokines (eotaxin, IP-10, MCP1, MIP1β, SCF, SDF-1α) and 8 growth factors (bNGF, BDNF, EGF, HGF, LIF, PIGF, VEGF-A, VEGF-D). Among them, FF MCP1, IFNγ, LIF and VEGF-D were significant lower in DHEA+ compared to DHEA- (*p*=0.03, 0.014, 0.031, 0.0161 respectively; **Figure 4**). No correlation was found between the significant cytokines and metabolites.

**Figure 4.**
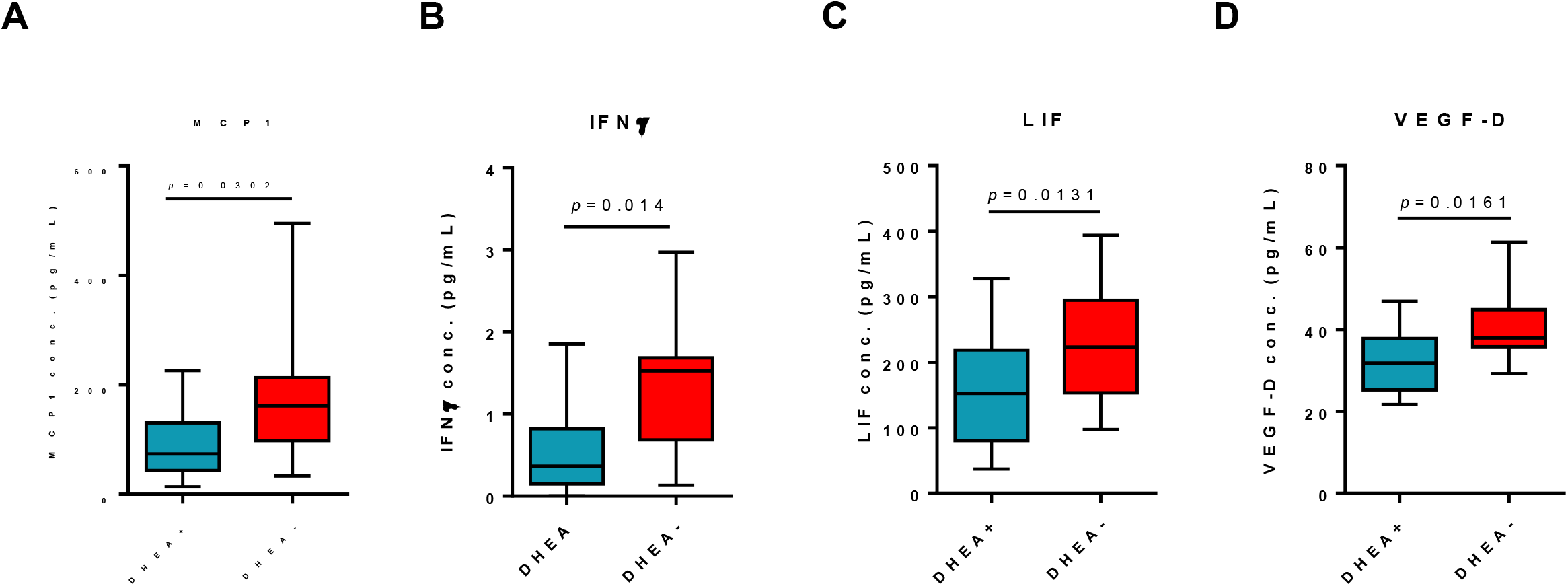
Significantly changed follicular fluid cytokines in DHEA+ and DHEA- patients. Among 45 cytokines, chemokines and growth factors measured by multiplex immunoassay, (A) MCP-1, (B) IFNγ, (C) LIF and (D) VEGF-D were significantly lower in poor ovarian response subjects with DHEA supplementation. Student’s t-tests were performed and p<0.05 is considered statistically significant.

### Clinical observations

Because of the treatment, DHEA-sulphate and free testosterone were significantly higher in the DHEA+ group compared to the DHEA- group (**Figure S4**). **Table 3** summarizes the study primary and secondary clinical outcomes. There were no significant differences for other of the matched criteria, consistent with other studies ^38–40^, although there were trends of increased retrieved oocytes (5.27□±□2.9 versus 4.Mtì]3.6) and metaphase II oocytes (4.0 ± 2.5 versus 3.5 ± 2.2) in DHEA+ group.

### Discussion

The follicular milieu provides oocytes with a specialized microenvironment that promotes the developmental competence of oocytes. It has been proposed that exogenous DHEA supplementation in POR patients with reduced follicle responsiveness to FSH, can optimize their response to ovarian stimulation for IVF among POR patients; however, the effect remains controversial.^4,13,41^ This LC-MS/MS metabolomics study extends the human FF metabolome space in terms of characterization of its constituents, providing new insights into the complexities of oocyte development especially in POR women,^42^ as well as with DHEA supplementation.^43, 44, 16–18^ Aside from previously reported constituents of FF such as linoleic acid,^16^ amino acids,^17^ and steroids including progesterone, testosterone^45^, this study also captured metabolic products of ovarian steroidogenesis, cholesterols and glucocorticoids, in the FF. Further, our FF metabolomics distinguished the DHEA+ group from the DHEA- group. This is despite the paucity of DHEA supplementation leading to clinical outcomes in this study, which was also observed by others,^38–40^ and that our identification of four significantly changed metabolites (glycerophosphocholine, linoleic acid, progesterone, and valine) which may alert us to the metabolic effects of exogenous DHEA supplementation.

Choline and derivatives are an emerging class of metabolites critical in developmental competence of fertilized oocytes,^18^ and glycerophosphorylcholine was found increased in the DHEA+ group. Glycerophosphorylcholine is formed from the breakdown of phosphatidylcholine, and is an organic osmolyte, plausibly affecting concentrations of other constitutes of FF,^15^ and regulation of the diffusion of compounds into FF necessary for folliculogenesis and oogenesis.^46^ PORs are known to exhibit a low diffusion of exogenous gonadotropin into FF, which is correlated with poor IVF outcomes.^47^ It is conceivable that DHEA induced the metabolism of phosphatidylcholine to glycerophosphorylcholine, and that a different DHEA supplementation regimen might lead to changes in FF composition, albeit at unknown effects on clinical outcomes. Progesterone is one of the key hormones for the progress of the first metiotic division in oocyte maturation, but changes to progesterone levels with DHEA supplementation has been fraught with controversies^48^. Our metabolomics study revealed for the first time that DHEA supplementation led to a decrease in FF progesterone levels, but to what impact lower progesterone induced by DHEA supplementation has on PORs remains to be investigated. Valine degradation has been previously reported in a iTRAQ proteomics study comparing competent versus incompetent buffalo oocyte proteome.^49^ In a metabolomics study, valine metabolism was also identified in bovine cumulus and cumulus-oocyte-complex-conditioned media that undergo oocyte maturation;^50^ although in both studies, valine was not directly detected in the omics profiling. In humans, degenerate oocytes or germinal vesicles that failed meiotically to reach metaphase II deplete valine more than competent oocytes, in other words lower valine levels in media which is consistent with our results, and suggest plausible biological roles of valine in oocyte maturation. Interestingly, we noted a segregation of DHEA+ patients with low and high level of valine, with the high valine group approaching concentrations of the DHEA- group. Together with valine’s high AUC value, it is tempting to speculate that valine might be a strong biomarker for monitoring individual DHEA supplementation. Linoleic acid is the most abundant polyunsaturated fatty acid in the bovine^51^ and human FF (**Figure 2**), and varying concentrations of linoleic acid have reportedly different effects on oocyte maturation. At a concentration of 100 μM, linoleic acid added to maturation media inhibits bovine oocyte maturation and subsequent blastocyst development through increasing prostaglandin E2 concentration in the medium, decreasing intracellular cAMP, decreasing phosphorylation of the MAPK1 and AKT and inhibited germinal vesicle breakdown.^51,52^ Conversely, at concentrations at 50 μM or below, linoleic acid improved oocyte quality by increasing the content of neutral lipids stored in lipid droplets.^52^ Our metabolomics results suggest that FF linoleic acid is another key biomarker for titrating and monitoring individual DHEA supplementation. Therefore, from published literature and our data suggest that DHEA might exert protective effects on oocytes in POR patients, albeit at an insufficient dose or treatment duration.

The elevated DHEA-sulphate levels coupled with a lack of difference in FF testosterone with DHEA supplementation suggest the following possibilities in POR patients: (i) inadequate DHEA conversion to testosterone due to polymorphism in *SULT2A1, CYP19A1 and FMR1* genes,^53^ or (ii) long CAG repeats in androgen receptor gene which is linked to its lower transcriptional activity at the promoters of genes involved in the metabolism of DHEA to testosterone.^54^ The former is unlikely: in a case-control study involving 94 subjects, androgen secretion was not impaired in pre-ovulatory follicles of POR compared to normal responders, and similar levels of follicular testosterone levels was reported.^55^ However, ethnicity and genetic predispositions might play a role as Chinese women are reported to have higher free androgens and African American women lower,^56^ which might explain their differences in pregnancy rates in association with IVF than those observed among other ethnic groups. Conversely, long CAG repeats is associated with risk of POR and oocyte insensitivity to androgenic stimulation,^57^ thus hinting a tenable rationale on the observed similar FF androgen levels between the DHEA- and DHEA+ subjects in this study. The abundance of cortisol in the DHEA+ subjects is an interesting finding, in particular that DHEA reduces circulating cortisol,^58^ indicating follicular versus systemic difference in how DHEA affects cortisol levels. *In vitro,* it was noted that DHEA suppresses cortisol activity,^59^ including the antagonist effects of DHEA on the anti-inflammatory responses induced by cortisol via glucocorticoid receptor-mediated pathways.^60^ It is noteworthy that high FF cortisol levels found in fertilized IVF individuals compared to unfertilized individuals led to the postulation that oocyte exposure to cortisol is required with oocyte maturation.^61^ The higher levels of FF cortisol observed in DHEA+ subjects therefore argues for a compensatory response to modulate the ratio of the two hormones to maintain for a favourable FF response to mature oocytes.^60^

Due to the highly confident identification based on MS/MS, and mass accuracy of LC-MS/MS-based metabolomics, we were able to distinguish progesterone from DHEA, an advantage over interference-prone immunoassays that face a cross-reactivity bioanalytical problem.^48^ Similarly, LC-MS/MS-based determination of androgens was preferred over immunoassays due to strong interference from DHEA.^62^ We did not detect E1 and E2; because for phenolic hydroxyl group of estrogens to act as proton donors, the signal would be more sensitive in the negative ion mode electrospray ionization^63^ than in the positive ion mode which was used in this study. Alternatively, derivatization with dansyl chloride, an amine-containing sulfonyl halide^64,65^ can be used for increased sensitivity but is more amenable to targeted LC-MS/MS.

In mouse models of polycystic ovary syndrome, treatment with DHEA resulted in increased production of cytokines such as serum TNFα, IL-6, IL12p70, and IFNγ.^66,67^ In this study, DHEA supplementation lead to the reduction of FF IFNγ, LIF, MCP-1, and VEGF-D levels. It appears that DHEA modulates chemokines and growth factors in POR FF without a clear Th1 or Th2 immune response as proposed.^40^ LIF or leukemia inhibitory factor is expressed in the ovary and controls follicular growth.^68^ It was reported that LIF suppressed the growth of primary, secondary, and early antral follicles in cultured ovarian tissues.^69^ The authors postulated that LIF produced in the late antral or graafian follicles is secreted to suppress the growth of the neighbouring primary, secondary, and early antral follicles as part of follicular growth.^15^ Interestingly, when hCG is administered in rhesus macaques, at 12 h follicular LIF levels increase and induce follicle rupture and ovulation and decrease at 24 h.^70^ In our study, the number of MII oocytes and oocytes trended higher in the DHEA+ group, suggesting that the biological roles of LIF might have been achieved (follicular maturation and rupture) but inadequate to generate a clinically significant outcome. *In vitro* results suggested that follicles produce VEGF-A, with VEGF-A inducing the expanding vasculature to support the increased needs growing follicles.^71^ The decrease in VEGF-A in DHEA+ individuals is intriguing. Fisher *et al.,* described that in cultured follicles, the rise in VEGF-A levels in faster-growing follicles are dependent on FSH dose and oxygen tension.^72^ There have been reports that DHEA inhibits oxygen consumption in neurons,^73^ tempting the postulation that DHEA inhibited oxygen consumption in follicle that subsequently led to lower production of VEGF-A in DHEA+ individuals. Further, the lack of correlation between the significant cytokines and metabolites suggest that DHEA converting to steroids which then modulate cytokine production within the follicular microenvironment is more complex than originally thought.

In conclusion, our study provided new insights to POR FF at the metabolome level, and as indicated from the FF metabolome analysis, exogenous DHEA to these patients altered the overall metabolome coverage and abundance to four metabolites. Hypotheses generated from this study included plausible mechanisms underlying DHEA metabolism, and the potential utility of glycerophosphocholine, linoleic acid, progesterone, and L-valine as markers to assess DHEA supplementation. Therefore, future directions include targeted quantitative LC-MS/MS approaches to be developed to detect and quantify four “responder” metabolites in approaches similar to those previously conducted on human peritoneal fluids and sera.^74–76^ to design treatment based on metabolomics profiles. Steroid hormones including testosterone should also be quantified via LC-MS/MS to establish baseline levels before commencing DHEA supplementation. Further, comparing POR and normal responders will provide further insights to the alteration of the FF metabolome, and reach a deeper understanding of underpinning pathophysiology to POR.

## Acknowledgements

This work was supported by SingHealth Foundation (SHF/CTG034/2010) to VVF, and the National Medical Research Council Centre Grant Programme (NMRC/CG/M003/2017) to YHL. JKYC received salary support from Singapore’s Ministry of Health’s National Medical Research Council (NMRC/CSA(SI)/008/2016).

## Authors contribution

VVF, HHT and JCKY recruited patients. JZ performed metabolomics and cytokine analyses. DB performed statistical analysis. YT provided expert help on metabolomics. LYH and JCKY supervised the work and resources. All authors agreed to submission of the manuscript and have agreed to the order of authors.

## Supplementary Figures

**Supplementary Figure 1.** Principle component analysis reveals DHEA+4 (arrow) as a potential outlier and was removed from subsequent analysis.

**Supplementary Figure 2.** (a) MS/MS spectra of pyridine at increasing eV. (b) Follicular fluid testerosterone levels as measured by metabolomics. DHEA+, POR subjects on DHEA supplementation and DHEA-without DHEA supplementation.

**Supplementary Figure 3.** (a) Dot Plots of Linoleic acid and L-Valine after removal of women with endometriosis (N=5), (b) ROC curves of Linoleic acid and L-Valine after removal of women with endometriosis (N=5).

**Supplementary Figure 4.** Histograms of estradiol, anti-müllerian hormone (AMH), DHEA-sulphate and insulin Growth Factor-1 (IGFBP-1) concentrations as determined by immunoassay. NS, not significant.

## Reference

(1) Surrey, E. S.; Schoolcraft, W. B. Evaluating Strategies for Improving Ovarian Response of the Poor Responder Undergoing Assisted Reproductive Techniques. Fertility and Sterility. 2000, pp 667–676. https://doi.org/10.1016/S0015-0282(99)00630-5.

(2) Devine, K.; Mumford, S. L.; Wu, M.; DeCherney, A. H.; Hill, M. J.; Propst, A. Diminished Ovarian Reserve in the United States Assisted Reproductive Technology Population: Diagnostic Trends among 181,536 Cycles from the Society for Assisted Reproductive Technology Clinic Outcomes Reporting System. In Fertility and Sterility; 2015; Vol. 104, pp 612–619.e3. https://doi.org/10.1016/j.fertnstert.2015.05.017.

(3) Ferraretti, A. P.; La Marca, A.; Fauser, B. C. J. M.; Tarlatzis, B.; Nargund, G.; Gianaroli, L.; ESHRE working group on Poor Ovarian Response Definition. ESHRE Consensus on the Definition of “poor Response” to Ovarian Stimulation for in Vitro Fertilization: The Bologna Criteria. Hum. Reprod. 2011, 26 (7), 1616–1624. https://doi.org/10.1093/humrep/der092.

(4) Kolibianakis, E. M.; Venetis, C. A.; Diedrich, K.; Tarlatzis, B. C.; Griesinger, G. Addition of Growth Hormone to Gonadotrophins in Ovarian Stimulation of Poor Responders Treated by In-Vitro Fertilization: A Systematic Review and Meta-Analysis. Human Reproduction Update. 2009, pp 613–622. https://doi.org/10.1093/humupd/dmp026.

(5) Haning, R. V; Flood, C. A.; Hackett, R. J.; Loughlin, J. S.; McClure, N.; Longcope, C. Metabolic Clearance Rate of Dehydroepiandrosterone Sulfate, Its Metabolism to Testosterone, and Its Intrafollicular Metabolism to Dehydroepiandrosterone, Androstenedione, Testosterone, and Dihydrotestosterone in Vivo. J. Clin. Endocrinol. Metab. 1991, 72 (5), 1088–1095. https://doi.org/10.1210/jcem-72-5-1088.

(6) Casson, P. R.; Santoro, N.; Elkind-Hirsch, K.; Carson, S. A.; Hornsby, P. J.; Abraham, G.; Buster, J. E. Postmenopausal Dehydroepiandrosterone Administration Increases Free Insulin-like Growth Factor-I and Decreases High-Density Lipoprotein: A Six-Month Trial. Fertil. Steril. 1998, 70 (1), 107–110. https://doi.org/10.1016/S0015-0282(98)00121-6.

(7) Barad, D.; Gleicher, N. Effect of Dehydroepiandrosterone on Oocyte and Embryo Yields, Embryo Grade and Cell Number in IVF. Hum. Reprod. 2006, 21 (11), 2845–2849. https://doi.org/10.1093/humrep/del254.

(8) Casson, P. R. Dehydroepiandrosterone Supplementation Augments Ovarian Stimulation in Poor Responders: A Case Series. Hum. Reprod. 2000, 15 (10), 2129–2132. https://doi.org/10.1093/humrep/15.10.2129.

(9) Burger, H. G. Androgen Production in Women. Fertil. Steril. 2002, 77 (4), S3–S5. https://doi.org/10.1016/S0015-0282(02)02985-0.

(10) Roy, S.; Mahesh, V. B.; Greenblatt, R. B. Effect of Dehydroepiandrosterone and Δ4-Androstenedione on the Reproductive Organs of Female Rats: Production of Cystic Changes in the Ovary. Nature 1962, 196 (4849), 42–43. https://doi.org/10.1038/196042a0.

(11) Broekmans, F. J.; Soules, M. R.; Fauser, B. C. Ovarian Aging: Mechanisms and Clinical Consequences. Endocr. Rev. 2009, 30 (5), 465–493. https://doi.org/10.1210/er.2009-0006.

(12) Barad, D.; Brill, H.; Gleicher, N. Update on the Use of Dehydroepiandrosterone Supplementation among Women with Diminished Ovarian Function. In Journal of Assisted Reproduction and Genetics; 2007; Vol. 24, pp 629–634. https://doi.org/10.1007/s10815-007-9178-x.

(13) Kyrou, D.; Kolibianakis, E. M.; Venetis, C. A.; Papanikolaou, E. G.; Bontis, J.; Tarlatzis, B. C. How to Improve the Probability of Pregnancy in Poor Responders Undergoing in Vitro Fertilization: A Systematic Review and Meta-Analysis. Fertil. Steril. 2009, 91 (3), 749–766. https://doi.org/10.1016/j.fertnstert.2007.12.077.

(14) Bächler, M.; Menshykau, D.; De Geyter, C.; Iber, D. Species-Specific Differences in Follicular Antral Sizes Result from Diffusion-Based Limitations on the Thickness of the Granulosa Cell Layer. MHR Basic Sci. Reprod. Med. 2014, 20 (3), 208–221. https://doi.org/10.1093/molehr/gat078.

(15) Rodgers, R. J.; Irving-Rodgers, H. F. Formation of the Ovarian Follicular Antrum and Follicular Fluid. Biol. Reprod. 2010, 82 (6), 1021–1029. https://doi.org/10.1095/biolreprod.109.082941.

(16) O’Gorman, A.; Wallace, M.; Cottell, E.; Gibney, M. J.; McAuliffe, F. M.; Wingfield, M.; Brennan, L. Metabolic Profiling of Human Follicular Fluid Identifies Potential Biomarkers of Oocyte Developmental Competence. Reproduction 2013, 146 (4), 389–395. https://doi.org/10.1530/REP-13-0184.

(17) Lan Xia; Zhao, X.; Sun, Y.; Hong, Y.; Yuping Gao, S. H. Metabolomic Profiling of Human Follicular Fluid from Patients with Repeated Failure of in Vitro Fertilization Using Gas Chromatography/Mass Spectrometry. Int. J. Clin. Experimenal Pathol. 2014, 7 (10), 7220–7229.

(18) Wallace, M.; Cottell, E.; Gibney, M. J.; McAuliffe, F. M.; Wingfield, M.; Brennan, L. An Investigation into the Relationship between the Metabolic Profile of Follicular Fluid, Oocyte Developmental Potential, and Implantation Outcome. Fertil. Steril. 2012, 97 (5), 1078–1084.e8. https://doi.org/10.1016/j.fertnstert.2012.01.122.

(19) Piñero-Sagredo, E.; Nunes, S.; de los Santos, M. J.; Celda, B.; Esteve, V. NMR Metabolic Profile of Human Follicular Fluid. NMR Biomed. 2010, 23 (5), 485–495. https://doi.org/10.1002/nbm.1488.

(20) Zamah, A. M.; Hassis, M. E.; Albertolle, M. E.; Williams, K. E. Proteomic Analysis of Human Follicular Fluid from Fertile Women. Clin. Proteomics 2015, 12 (1), 5. https://doi.org/10.1186/s12014-015-9077-6.

(21) Angelucci, S.; Ciavardelli, D.; Di Giuseppe, F.; Eleuterio, E.; Sulpizio, M.; Tiboni, G. M.; Giampietro, F.; Palumbo, P.; Di Ilio, C. Proteome Analysis of Human Follicular Fluid. Biochim. Biophys. Acta - Proteins Proteomics 2006, 1764 (11), 1775–1785. https://doi.org/10.1016/j.bbapap.2006.09.001.

(22) Jarkovska, K.; Martinkova, J.; Liskova, L.; Halada, P.; Moos, J.; Rezabek, K.; Gadher, S. J.; Kovarova, H. Proteome Mining of Human Follicular Fluid Reveals a Crucial Role of Complement Cascade and Key Biological Pathways in Women Undergoing in Vitro Fertilization. J. Proteome Res. 2010, 9 (3), 1289–1301. https://doi.org/10.1021/pr900802u.

(23) Hanrieder, J.; Nyakas, A.; Naessén, T.; Bergquist, J. Proteomic Analysis of Human Follicular Fluid Using an Alternative Bottom-Up Approach. J. Proteome Res. 2008, 7 (1), 443–449. https://doi.org/10.1021/pr070277z.

(24) Estes, S. J.; Ye, B.; Qiu, W.; Cramer, D.; Hornstein, M. D.; Missmer, S. A. A Proteomic Analysis of IVF Follicular Fluid in Women ≤32 Years Old. Fertil. Steril. 2009, 92 (5), 1569–1578. https://doi.org/10.1016/j.fertnstert.2008.08.120.

(25) Chen, F.; Spiessens, C.; D’Hooghe, T.; Peeraer, K.; Carpentier, S. Follicular Fluid Biomarkers for Human in Vitro Fertilization Outcome: Proof of Principle. Proteome Sci. 2016, 14 (1), 17. https://doi.org/10.1186/s12953-016-0106-9.

(26) Yakin, K.; Urman, B. DHEA as a Miracle Drug in the Treatment of Poor Responders; Hype or Hope? Hum. Reprod. 2011, 26 (8), 1941–1944. https://doi.org/10.1093/humrep/der150.

(27) Sönmezer, M.; Özmen, B.; Çil, A. P.; Özkavukçu, S.; Taşçi, T.; Olmuş, H.; Atabeko□lu, C. S. Dehydroepiandrosterone Supplementation Improves Ovarian Response and Cycle Outcome in Poor Responders. Reprod. Biomed. Online 2009, 19 (4), 508–513. https://doi.org/10.1016/j.rbmo.2009.06.006.

(28) Cui, L.; Lu, H.; Lee, Y. H. Challenges and Emergent Solutions for LC-MS/MS Based Untargeted Metabolomics in Diseases. Mass Spectrom. Rev. 2018. https://doi.org/10.1002/mas.21562.

(29) Hazeldine, J.; Arlt, W.; Lord, J. M. Dehydroepiandrosterone as a Regulator of Immune Cell Function. J. Steroid Biochem. Mol. Biol. 2010, 120 (2–3), 127–136. https://doi.org/10.1016/j.jsbmb.2009.12.016.

(30) Mason, H. D.; Margara, R.; Winston, R. M.; Seppala, M.; Koistinen, R.; Franks, S. Insulin-like Growth Factor-I (IGF-I) Inhibits Production of IGF-Binding Protein-1 While Stimulating Estradiol Secretion in Granulosa Cells from Normal and Polycystic Human Ovaries. J. Clin. Endocrinol. Metab. 1993, 76 (5), 1275–1279. https://doi.org/10.1210/jcem.76.5.7684393.

(31) Lee, Y. H.; Goh, W. W. Bin; Ng, C. K.; Raida, M.; Wong, L.; Lin, Q.; Boelsterli, U. a; Chung, M. C. M. Integrative Toxicoproteomics Implicates Impaired Mitochondrial Glutathione Import as an Off-Target Effect of Troglitazone. J. Proteome Res. 2013, 12 (6), 2933–2945. https://doi.org/10.1021/pr400219s.

(32) Cui, L.; Fang, J.; Ooi, E. E.; Lee, Y. H. Serial Metabolome Changes in a Prospective Cohort of Subjects with Influenza Viral Infection and Comparison with Dengue Fever. J. Proteome Res. 2017, 16 (7), 2614–2622. https://doi.org/10.1021/acs.jproteome.7b00173.

(33) Peter Durairaj, R. R.; Aberkane, A.; Polanski, L.; Maruyama, Y.; Baumgarten, M.; Lucas, E. S.; Quenby, S.; Chan, J. K. Y.; Raine-Fenning, N.; Brosens, J. J.; Van de Velde, H.; Lee, Y. H. Deregulation of the Endometrial Stromal Cell Secretome Precedes Embryo Implantation Failure. MHR Basic Sci. Reprod. Med. 2017, 1–10. https://doi.org/10.1093/molehr/gax023.

(34) Wishart, D. S.; Tzur, D.; Knox, C.; Eisner, R.; Guo, A. C.; Young, N.; Cheng, D.; Jewell, K.; Arndt, D.; Sawhney, S.; Fung, C.; Nikolai, L.; Lewis, M.; Coutouly, M.-A.; Forsythe, I.; Tang, P.; Shrivastava, S.; Jeroncic, K.; Stothard, P.; Amegbey, G.; Block, D.; Hau, D. D.; Wagner, J.; Miniaci, J.; Clements, M.; Gebremedhin, M.; Guo, N.; Zhang, Y.; Duggan, G. E.; Macinnis, G. D.; Weljie, A. M.; Dowlatabadi, R.; Bamforth, F.; Clive, D.; Greiner, R.; Li, L.; Marrie, T.; Sykes, B. D.; Vogel, H. J.; Querengesser, L. HMDB: The Human Metabolome Database. Nucleic Acids Res. 2007, 35 (Database issue), D521–6. https://doi.org/10.1093/nar/gkl923.

(35) Williams, J. R.; Boehm, J. C. Studies on the Synthesis of Dehydroepiandrosterone (DHEA) Phosphatide. Steroids 1995, 60 (4), 333–336.

(36) Nielsen, M. E.; Rasmussen, I. A.; Kristensen, S. G.; Christensen, S. T.; Mollgard, K.; Wreford Andersen, E.; Byskov, A. G.; Yding Andersen, C. In Human Granulosa Cells from Small Antral Follicles, Androgen Receptor MRNA and Androgen Levels in Follicular Fluid Correlate with FSH Receptor MRNA. Mol. Hum. Reprod. 2011, 17 (1), 63–70. https://doi.org/10.1093/molehr/gaq073.

(37) Gleicher, N.; Weghofer, A.; Barad, D. H. The Role of Androgens in Follicle Maturation and Ovulation Induction: Friend or Foe of Infertility Treatment? Reprod. Biol. Endocrinol. 2011, 9 (1), 116. https://doi.org/10.1186/1477-7827-9-116.

(38) Wiser, A.; Gonen, O.; Ghetler, Y.; Shavit, T.; Berkovitz, A.; Shulman, A. Addition of Dehydroepiandrosterone (DHEA) for Poor-Responder Patients before and during IVF Treatment Improves the Pregnancy Rate: A Randomized Prospective Study. Hum. Reprod. 2010, 25 (10), 2496–2500. https://doi.org/10.1093/humrep/deq220.

(39) Hu, Q.; Hong, L.; Nie, M.; Wang, Q.; Fang, Y.; Dai, Y.; Zhai, Y.; Wang, S.; Yin, C.; Yang, X. The Effect of Dehydroepiandrosterone Supplementation on Ovarian Response Is Associated with Androgen Receptor in Diminished Ovarian Reserve Women. J. Ovarian Res. 2017, 10 (1), 32. https://doi.org/10.1186/s13048-017-0326-3.

(40) Zhang, J.; Qiu, X.; Gui, Y.; Xu, Y.; Li, D.; Wang, L. Dehydroepiandrosterone Improves the Ovarian Reserve of Women with Diminished Ovarian Reserve and Is a Potential Regulator of the Immune Response in the Ovaries. Biosci. Trends 2015, 9 (6), 350–359. https://doi.org/10.5582/bst.2015.01154.

(41) Duffy, J. M.; Ahmad, G.; Mohiyiddeen, L.; Nardo, L. G.; Watson, A. Growth Hormone for in Vitro Fertilization. Cochrane database Syst. Rev. 2010, No. 1, CD000099. https://doi.org/10.1002/14651858.CD000099.pub3.

(42) Emori, M. M.; Drapkin, R. The Hormonal Composition of Follicular Fluid and Its Implications for Ovarian Cancer Pathogenesis. Reprod. Biol. Endocrinol. 2014, 12 (1), 60. https://doi.org/10.1186/1477-7827-12-60.

(43) Bedaiwy, M.; Shahin, A. Y.; AbulHassan, A. M.; Goldberg, J. M.; Sharma, R. K.; Agarwal, A.; Falcone, T. Differential Expression of Follicular Fluid Cytokines: Relationship to Subsequent Pregnancy in IVF Cycles. Reprod. Biomed. Online 2007, 15 (3), 321–325. https://doi.org/10.1016/S1472-6483(10)60346-X.

(44) Baskind, N. E.; Orsi, N. M.; Sharma, V. Follicular-Phase Ovarian Follicular Fluid and Plasma Cytokine Profiling of Natural Cycle in Vitro Fertilization Patients. Fertil. Steril. 2014, 102 (2), 410–418. https://doi.org/10.1016/j.fertnstert.2014.04.032.

(45) Wen, X.; Li, D.; Tozer, A. J.; Docherty, S. M.; Iles, R. K. Estradiol, Progesterone, Testosterone Profiles in Human Follicular Fluid and Cultured Granulosa Cells from Luteinized Pre-Ovulatory Follicles. Reprod. Biol. Endocrinol. 2010, 8 (1), 117. https://doi.org/10.1186/1477-7827-8-117.

(46) Hsueh, A. J. W.; Kawamura, K.; Cheng, Y.; Fauser, B. C. J. M. Intraovarian Control of Early Folliculogenesis. Endocr. Rev. 2015, 36 (1), 1–24. https://doi.org/10.1210/er.2014-1020.

(47) Nagata, Y.; Honjou, K.; Sonoda, M.; Sumii, Y.; Inoue, Y.; Kawarabayashi, T. Pharmacokinetics of Exogenous Gonadotropin and Ovarian Response in in Vitro Fertilization. Fertil. Steril. 1999, 72 (2), 235–239. https://doi.org/10.1016/S0015-0282(99)00228-9.

(48) Forman, E. J.; Franasiak, J. M.; Scott, R. T. Elevated Progesterone Levels in Women on DHEA Supplementation Likely Represent Assay Interference. J. Assist. Reprod. Genet. 2015, 32 (4), 661–661. https://doi.org/10.1007/s10815-015-0442-1.

(49) Chen, L.; Zhai, L.; Qu, C.; Zhang, C.; Li, S.; Wu, F.; Qi, Y.; Lu, F.; Xu, P.; Li, X.; Shi, D. Comparative Proteomic Analysis of Buffalo Oocytes Matured in Vitro Using ITRAQ Technique. Sci. Rep. 2016, 6 (1), 31795. https://doi.org/10.1038/srep31795.

(50) Uhde, K.; van Tol, H. T. A.; Stout, T. A. E.; Roelen, B. A. J. Metabolomic Profiles of Bovine Cumulus Cells and Cumulus-Oocyte-Complex-Conditioned Medium during Maturation in Vitro. Sci. Rep. 2018, 8 (1), 9477. https://doi.org/10.1038/s41598-018-27829-9.

(51) Marei, W. F.; Wathes, D. C.; Fouladi-Nashta, A. A. Impact of Linoleic Acid on Bovine Oocyte Maturation and Embryo Development. Reproduction 2010, 139 (6), 979–988. https://doi.org/10.1530/REP-09-0503.

(52) Carro, M.; Buschiazzo, J.; Ríos, G. L.; Oresti, G. M.; Alberio, R. H. Linoleic Acid Stimulates Neutral Lipid Accumulation in Lipid Droplets of Maturing Bovine Oocytes. Theriogenology 2013, 79 (4), 687–694. https://doi.org/10.1016/j.theriogenology.2012.11.025.

(53) Shohat-Tal, A.; Sen, A.; Barad, D. H.; Kushnir, V.; Gleicher, N. Genetics of Androgen Metabolism in Women with Infertility and Hypoandrogenism. Nat. Rev. Endocrinol. 2015, 11 (7), 429–441. https://doi.org/10.1038/nrendo.2015.64.

(54) Borgbo, T.; Macek, M.; Chrudimska, J.; Jeppesen, J. V.; Hansen, L. L.; Andersen, C. Y. Size Matters: Associations between the Androgen Receptor CAG Repeat Length and the Intrafollicular Hormone Milieu. Mol. Cell. Endocrinol. 2016, 419, 12–17. https://doi.org/10.1016/j.mce.2015.09.015.

(55) De Los Santos, M. J.; García-Laez, V.; Beltrán, D.; Labarta, E.; Zuzuarregui, J. L.; Alamá, P.; Gámiz, P.; Crespo, J.; Bosch, E.; Pellicer, A. The Follicular Hormonal Profile in Low-Responder Patients Undergoing Unstimulated Cycles: Is It Hypoandrogenic? Hum. Reprod. 2013, 28 (1), 224–229. https://doi.org/10.1093/humrep/des349.

(56) Sutton-Tyrrell, K. Sex Hormone-Binding Globulin and the Free Androgen Index Are Related to Cardiovascular Risk Factors in Multiethnic Premenopausal and Perimenopausal Women Enrolled in the Study of Women Across the Nation (SWAN). Circulation 2005, 111 (10), 1242–1249. https://doi.org/10.1161/01.CIR.0000157697.54255.CE.

(57) Lledo, B.; Llácer, J.; Ortiz, J. A.; Martinez, B.; Morales, R.; Bernabeu, R. A Pharmacogenetic Approach to Improve Low Ovarian Response: The Role of CAG Repeats Length in the Androgen Receptor Gene. Eur. J. Obstet. Gynecol. Reprod. Biol. 2018, 227, 41–45. https://doi.org/10.1016/j.ejogrb.2018.06.001.

(58) Kroboth, P. D.; Amico, J. A.; Stone, R. A.; Folan, M.; Frye, R. F.; Kroboth, F. J.; Bigos, K. L.; Fabian, T. J.; Linares, A. M.; Pollock, B. G.; Hakala, C. Influence of DHEA Administration on 24-Hour Cortisol Concentrations. J. Clin. Psychopharmacol. 2003, 23 (1), 96–99. https://doi.org/10.1097/00004714-200302000-00014.

(59) Buoso, E.; Lanni, C.; Molteni, E.; Rousset, F.; Corsini, E.; Racchi, M. Opposing Effects of Cortisol and Dehydroepiandrosterone on the Expression of the Receptor for Activated C Kinase 1: Implications in Immunosenescence. Exp. Gerontol. 2011, 46 (11), 877–883. https://doi.org/10.1016/j.exger.2011.07.007.

(60) Kamin, H. S.; Kertes, D. A. Cortisol and DHEA in Development and Psychopathology. Horm. Behav. 2017, 89, 69–85. https://doi.org/10.1016/j.yhbeh.2016.11.018.

(61) Keay, S. D. Higher Cortisol:Cortisone Ratios in the Preovulatory Follicle of Completely Unstimulated IVF Cycles Indicate Oocytes with Increased Pregnancy Potential. Hum. Reprod. 2002, 17 (9), 2410–2414. https://doi.org/10.1093/humrep/17.9.2410.

(62) Heald, A. H.; Butterworth, A.; Kane, J. W.; Borzomato, J.; Taylor, N. F.; Layton, T.; Kilpatrick, E. S.; Rudenski, A. Investigation into Possible Causes of Interference in Serum Testosterone Measurement in Women. Ann. Clin. Biochem. 2006, 43 (3), 189–195. https://doi.org/10.1258/000456306776865106.

(63) Guo, T.; Gu, J.; Soldin, O. P.; Singh, R. J.; Soldin, S. J. Rapid Measurement of Estrogens and Their Metabolites in Human Serum by Liquid Chromatography-Tandem Mass Spectrometry without Derivatization. Clin. Biochem. 2008, 41 (9), 736–741. https://doi.org/10.1016/j.clinbiochem.2008.02.009.

(64) Nelson, R. E. Liquid Chromatography-Tandem Mass Spectrometry Assay for Simultaneous Measurement of Estradiol and Estrone in Human Plasma. Clin. Chem. 2004, 50 (2), 373–384. https://doi.org/10.1373/clinchem.2003.025478.

(65) Anari, M. R.; Bakhtiar, R.; Zhu, B.; Huskey, S.; Franklin, R. B.; Evans, D. C. Derivatization of Ethinylestradiol with Dansyl Chloride To Enhance Electrospray Ionization: Application in Trace Analysis of Ethinylestradiol in Rhesus Monkey Plasma. Anal. Chem. 2002, 74 (16), 4136–4144. https://doi.org/10.1021/ac025712h.

(66) Sander, V.; Luchetti, C. G.; Solano, M. E.; Elia, E.; Di Girolamo, G.; Gonzalez, C.; Motta, A. B. Role of the N, N’-Dimethylbiguanide Metformin in the Treatment of Female Prepuberal BALB/c Mice Hyperandrogenized with Dehydroepiandrosterone. Reproduction 2006, 131 (3), 591–602. https://doi.org/10.1530/rep.1.00941.

(67) Solano, M. E.; Sander, V. A.; Ho, H.; Motta, A. B.; Arck, P. C. Systemic Inflammation, Cellular Influx and up-Regulation of Ovarian VCAM-1 Expression in a Mouse Model of Polycystic Ovary Syndrome (PCOS). J. Reprod. Immunol. 2011, 92 (1–2), 33–44. https://doi.org/10.1016/j.jri.2011.09.003.

(68) Abir, R.; Fisch, B.; Jin, S.; Barnnet, M.; Freimann, S.; Van den Hurk, R.; Feldberg, D.; Nitke, S.; Krissi, H.; Ao, A. Immunocytochemical Detection and RT-PCR Expression of Leukaemia Inhibitory Factor and Its Receptor in Human Fetal and Adult Ovaries. Mol. Hum. Reprod. 2004. https://doi.org/10.1093/molehr/gah047.

(69) Komatsu, K.; Koya, T.; Wang, J.; Yamashita, M.; Kikkawa, F.; Iwase, A. Analysis of the Effect of Leukemia Inhibitory Factor on Follicular Growth in Cultured Murine Ovarian Tissue. Biol. Reprod. 2015, 93 (1). https://doi.org/10.1095/biolreprod.115.128421.

(70) Murphy, M. J.; Halow, N. G.; Royer, P. A.; Hennebold, J. D. Leukemia Inhibitory Factor Is Necessary for Ovulation in Female Rhesus Macaques. Endocrinology 2016, 157 (11), 4378–4387. https://doi.org/10.1210/en.2016-1283.

(71) Xu, J.; Bernuci, M. P.; Lawson, M. S.; Yeoman, R. R.; Fisher, T. E.; Zelinski, M. B.; Stouffer, R. L. Survival, Growth, and Maturation of Secondary Follicles from Prepubertal, Young, and Older Adult Rhesus Monkeys during Encapsulated Three-Dimensional Culture: Effects of Gonadotropins and Insulin. Reproduction 2010. https://doi.org/10.1530/REP-10-0284.

(72) Fisher, T. E.; Molskness, T. A.; Villeda, A.; Zelinski, M. B.; Stouffer, R. L.; Xu, J. Vascular Endothelial Growth Factor and Angiopoietin Production by Primate Follicles during Culture Is a Function of Growth Rate, Gonadotrophin Exposure and Oxygen Milieu. Hum. Reprod. 2013, 28 (12), 3263–3270. https://doi.org/10.1093/humrep/det337.

(73) Safiulina, D.; Peet, N.; Seppet, E.; Zharkovsky, A.; Kaasik, A. Dehydroepiandrosterone Inhibits Complex I of the Mitochondrial Respiratory Chain and Is Neurotoxic In Vitro and In Vivo at High Concentrations. Toxicol. Sci. 2006, 93 (2), 348–356. https://doi.org/10.1093/toxsci/kfl064.

(74) Lee, Y. H.; Yang, J. X.; Allen, J. C.; Tan, C. S.; Chern, B. S. M.; Tan, T. Y.; Tan, H. H.; Mattar, C. N. Z.; Chan, J. K. Y. Elevated Peritoneal Fluid Ceramides in Human Endometriosis-Associated Infertility and Their Effects on Mouse Oocyte Maturation. Fertil. Steril. 2018. https://doi.org/10.1016/j.fertnstert.2018.05.003.

(75) Lee, Y. H.; Tan, C. W.; Venkatratnam, A.; Tan, C. S.; Cui, L.; Loh, S. F.; Griffith, L.; Tannenbaum, S. R.; Chan, J. K. Y. Dysregulated Sphingolipid Metabolism in Endometriosis. J. Clin. Endocrinol. Metab. 2014, 99 (10), E1913–21. https://doi.org/10.1210/jc.2014-1340.

(76) Lee, Y. H.; Cui, L.; Fang, J.; Chern, B. S. M.; Tan, H. H.; Chan, J. K. Y. Limited Value of Pro-Inflammatory Oxylipins and Cytokines as Circulating Biomarkers in Endometriosis – a Targeted ‘omics Study. Sci. Rep. 2016, 6, 26117. https://doi.org/10.1038/srep26117.

