## Supplementary Figures for "Follicular fluid metabolome and cytokinome profiles in poor ovarian responders and the impact of dehydroepiandrosterone supplementation"

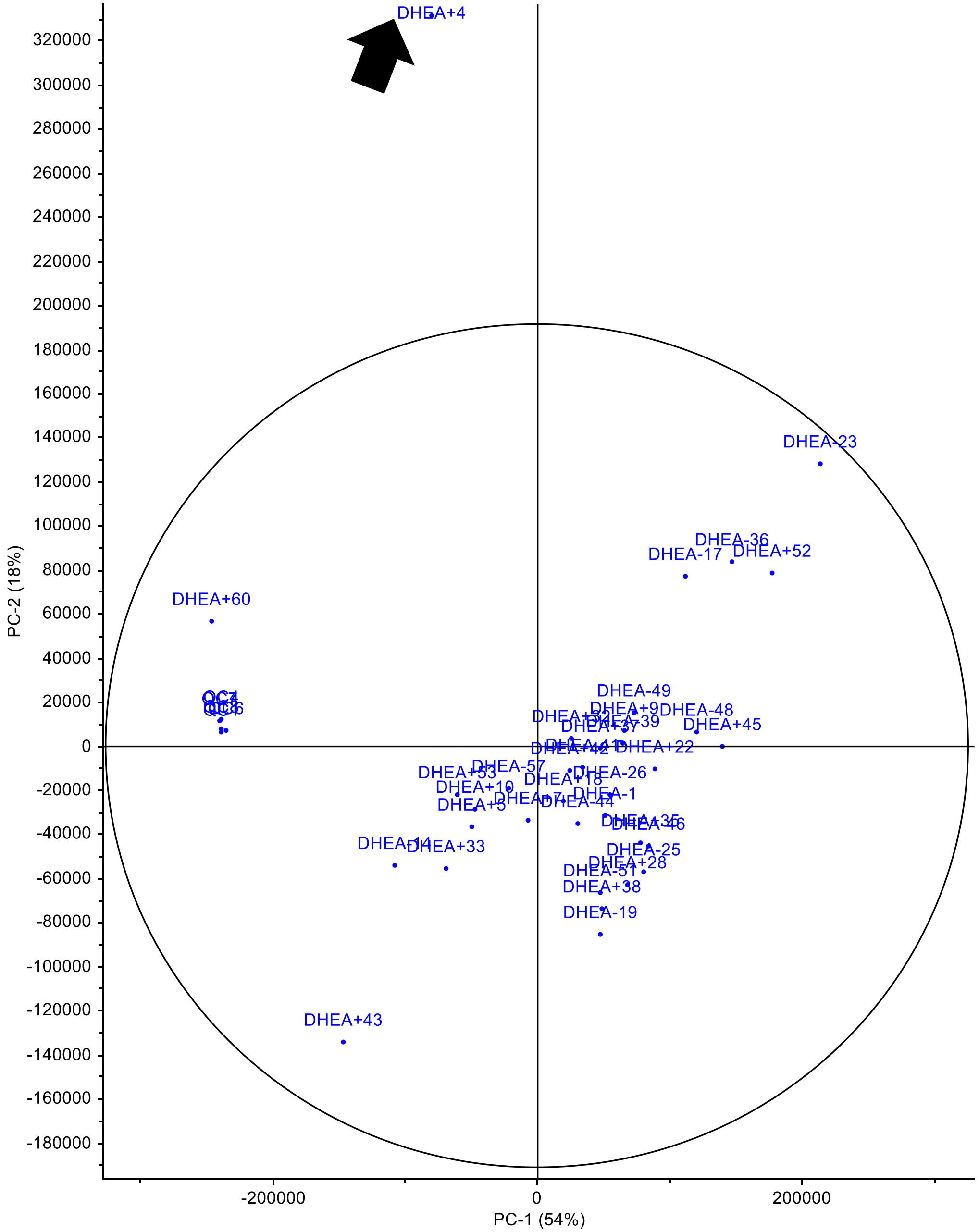


**Figure S1.** Principle component analysis reveals DHEA+4 (arrow) as a potential outlier and was removed from subsequent analysis.

**a**

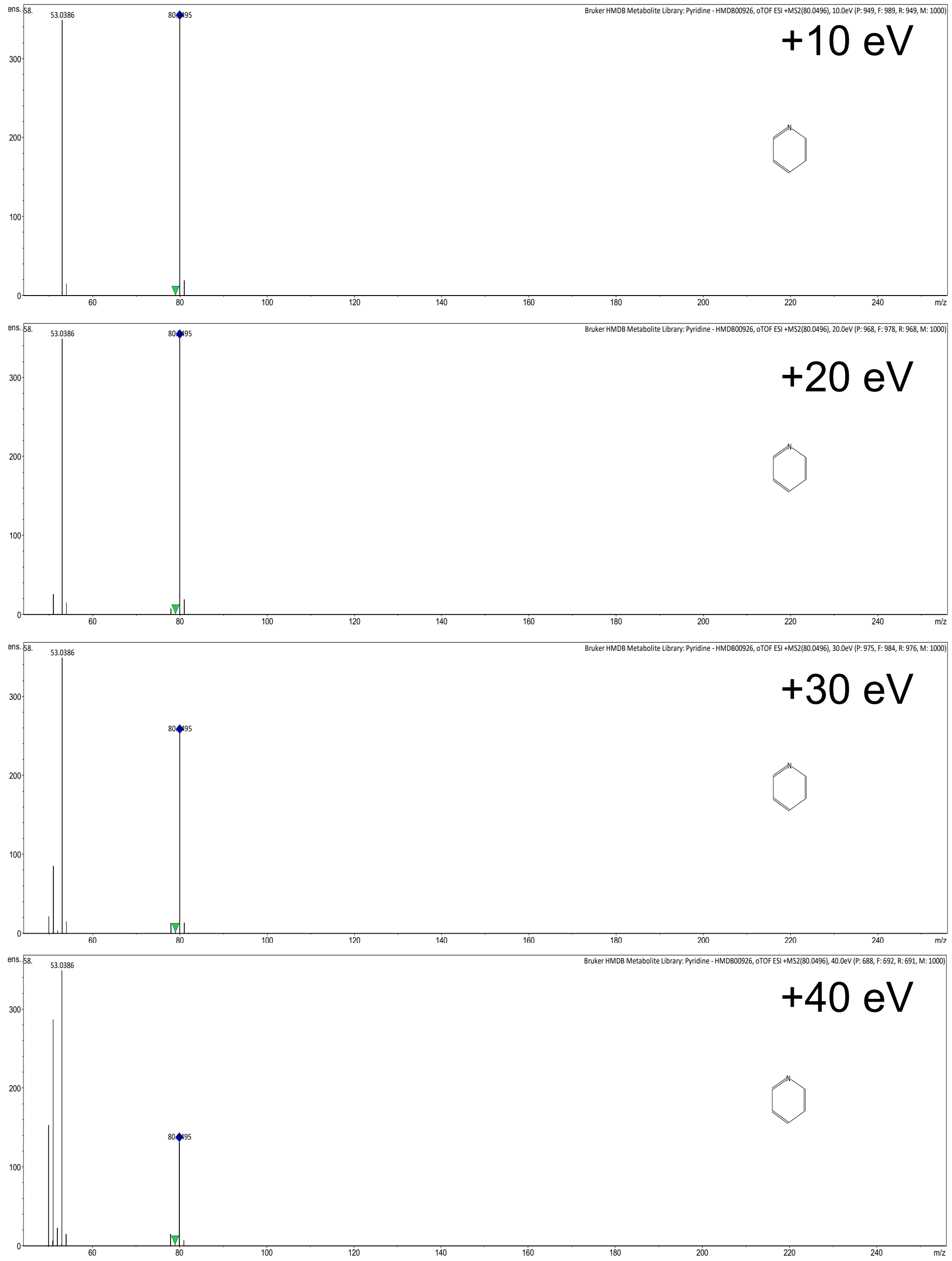


**b**

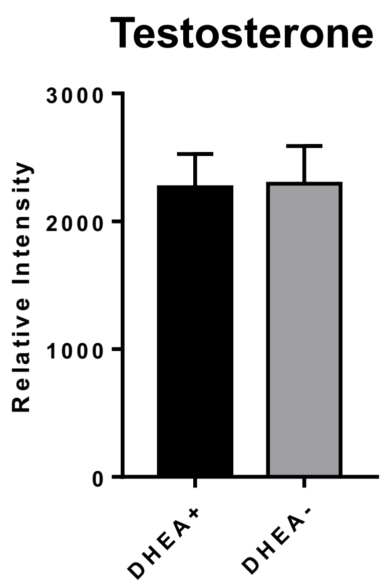


**Figure S2.** (a) MS/MS spectra of pyridine at increasing eV. (b) Follicular fluid testerosterone levels as measured by metabolomics. DHEA+, POR subjects on DHEA supplementation and DHEA- without DHEA supplementation.

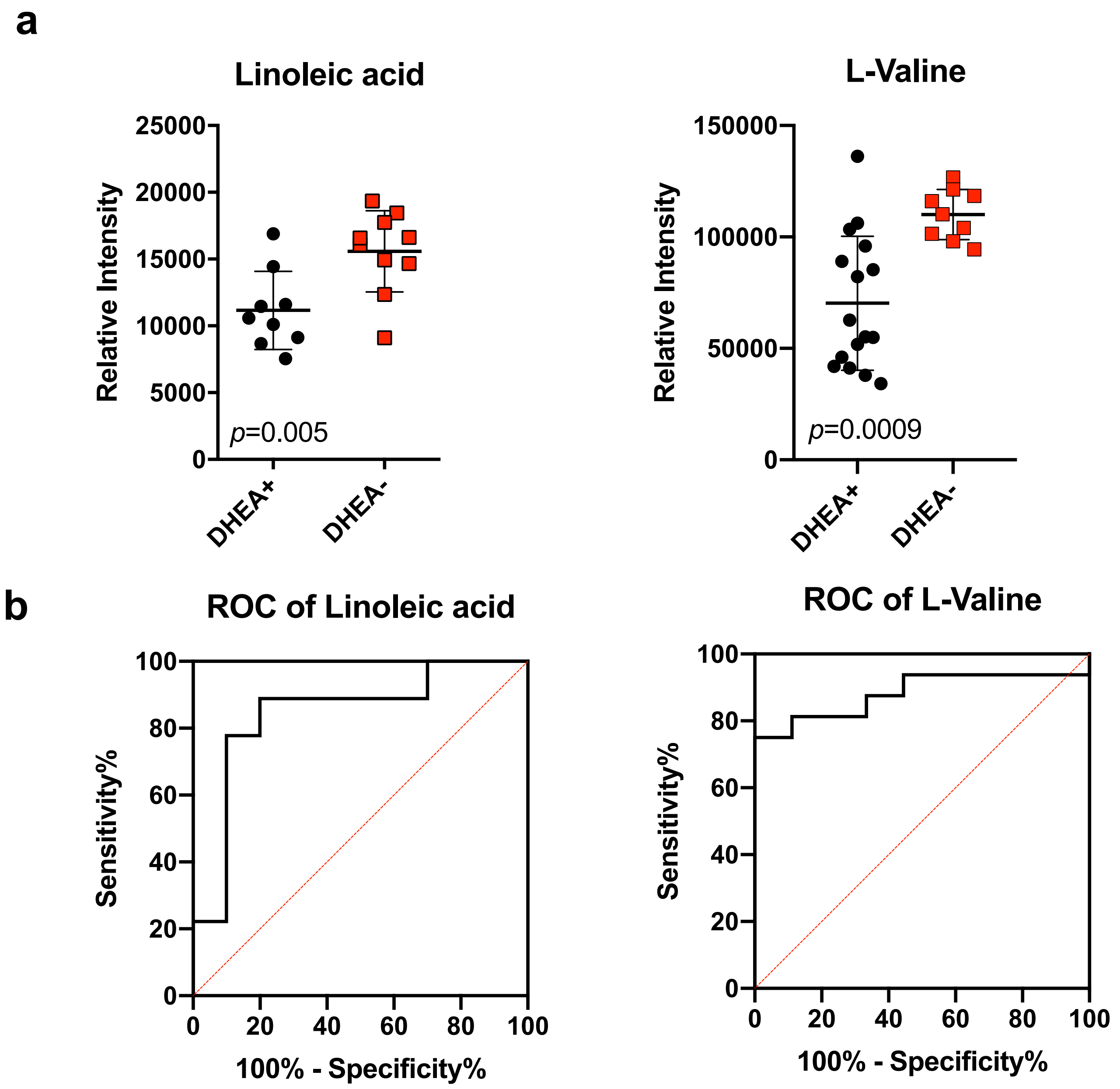


**Figure S3.** (a) Dot Plots of Linoleic acid and L-Valine after removal of women with endometriosis (*N*=5), (b) ROC curves of Linoleic acid and L-Valine after removal of women with endometriosis (*N*=5).

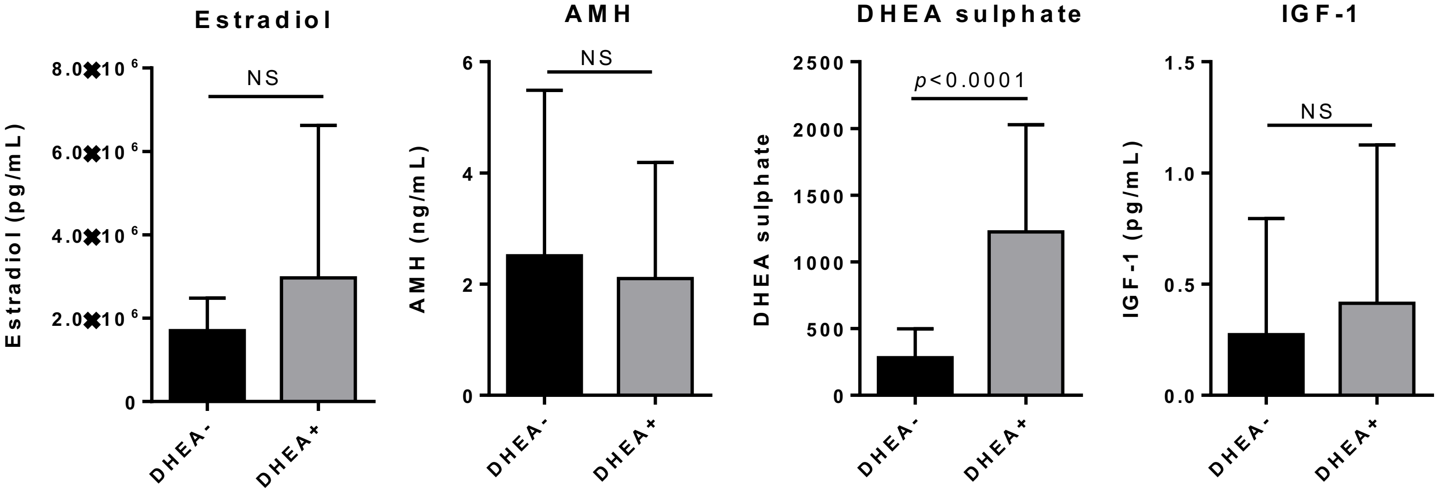


**Figure S4.** Histograms of estradiol, anti-müllerian hormone (AMH), DHEA-sulphate and insulin Growth Factor-1 (IGFBP-1) concentrations as determined by immunoassay. NS, not significant.
